# Genetic interactions derived from high-throughput phenotyping of 7,350 yeast cell cycle mutants

**DOI:** 10.1101/785840

**Authors:** Jenna E. Gallegos, Neil R. Adames, Mark F. Rogers, Pavel Kraikivski, Aubrey Ibele, Kevin Nurzynski-Loth, Eric Kudlow, T.M. Murali, John J. Tyson, Jean Peccoud

## Abstract

Over the last 30 years, computational biologists have developed increasingly realistic mathematical models of the regulatory networks controlling the division of eukaryotic cells. These models capture data resulting from two complementary experimental approaches: low-throughput experiments aimed at extensively characterizing the functions of small numbers of genes, and large-scale genetic interaction screens that provide a systems-level perspective on the cell division process. The former is insufficient to capture the interconnectivity of the genetic control network, while the latter is fraught with irreproducibility issues. Here, we describe a hybrid approach in which the genetic interactions between 36 cell-cycle genes are quantitatively estimated by high-throughput phenotyping with an unprecedented number of biological replicates. Using this approach, we identify a subset of high-confidence genetic interactions, which we use to refine a previously published mathematical model of the cell cycle. We also present a quantitative dataset of the growth rate of these mutants under six different media conditions in order to inform future cell cycle models.

**Author Summary:** The process of cell division, also called the cell cycle, is controlled by a highly complex network of interconnected genes. If this process goes awry, diseases such as cancer can result. In order to unravel the complex interactions within the cell cycle control network, computational biologists have developed mathematical models that describe how different cell cycle genes are related. These models are built using large datasets describing the effect of mutating one or more genes within the network. In this manuscript, we present a novel method for producing such datasets. Using our method, we generate 7,350 yeast mutants to explore the interactions between key cell cycle genes. We measure the effect of the mutations by monitoring the growth rate of the yeast mutants under different environmental conditions. We use our mutants to revise an existing model of the yeast cell cycle and present a dataset of ∼44,000 gene by environment combinations as a resource to the yeast genetics and modeling communities.

## Introduction

Eukaryotic cells grow and divide using a highly conserved and integrated network of positive and negative controls that ensure genomic integrity and maintain cell size within reasonable bounds. Proper control of the cell division cycle is essential for competitive fitness, embryonic development and maturation, and tissue homeostasis. Failure in these control mechanisms may result in cell death, developmental defects, tissue dysplasia, or cancers. One of the foremost model organisms for unraveling the molecular mechanisms of cell cycle control is the budding yeast *Saccharomyces cerevisiae*. Several hundred yeast mutants, generated in dozens of research laboratories over the past 40 years, have led to the discovery and characterization of many genes and proteins that regulate progression through the cell cycle^1^. Because of the intense labor involved in these experiments, individual laboratories have tended to focus on small numbers of genes and proteins involved in sub-sections of the extensive network of gene/protein interactions that control cell cycle events. This reductionist approach was necessary in the early stages of identifying and characterizing the molecular regulatory system, but it carries with it the danger of missing higher levels of network organization and their phenotypic consequences ^2–4^.

In contrast to a detailed, reductionist experimental approach, which builds a regulatory network from the bottom up, a systems-level approach seeks to provide a more global and less biased view of regulatory networks. Systems biologists can uncover key regulatory interactions and network architectures that bottom-up practitioners may have missed ^5, 6^. Unfortunately, the top-down, pan-genome approach, while good for generating hypotheses, is usually poor for testing hypotheses because the experiments are mostly correlative, and the data is often plagued by problems of accuracy and reproducibility. Combining a variety of ‘omics’ studies may help to overcome these challenges, but it is often difficult to integrate disparate data sets into a single network model ^7–10^. Ideally, one should combine top-down and bottom-up data, but huge discrepancies of scale between these two data types present barriers to integrating and understanding the hypotheses derived from each approach ^11–17^.

To mitigate these problems, many researchers, including ourselves, have developed detailed mathematical models that integrate top-down and bottom-up approaches in order to describe the molecular mechanisms that underlie cell cycle regulation in budding yeast ^4, 17–22^. The governing equations of the model are simulated on a computer, and the model (the ‘wiring diagram’ of molecular interactions) is adjusted until it generates dynamic behaviors that reflect the documented molecular changes and general network behaviors observed in cells (e.g., cell viability, timing of cell cycle events, cell size at birth, response to DNA damage or chromosome misalignment at mitosis) ^23–26^. Often, the documented data is missing detailed molecular information, such as protein concentrations and rate constants of crucial reactions, but fitting the model (i.e., fine-tuning the parameter values) to extensive sets of phenotypic data usually introduces strong constraints on these unknown parameters ^19, 27^. In this way, mathematical models can refine our understanding of the molecular mechanisms underlying cell cycle progression and test if the proposed network architecture and kinetic rate-constant estimates are consistent with both bottom-up and top-down observations.

One of the major problems when developing large mathematical models of the cell cycle has been the lack of consistent data sets. It has been challenging to compare data collected on cell cycle mutants that often have different genetic backgrounds, whose phenotypes are usually descriptive rather than quantitative, and whose phenotypes are assessed under inconsistent conditions. These problems leave the modeler with the difficult task of curating, interpreting and consolidating inconsistent and sometimes unreliable experimental results.

A particularly pernicious example of this problem is the use of the ‘synthetic lethal’ (SL) phenotype of double-mutant yeast cells in the development and calibration of mathematical models of the budding yeast cell cycle. Synthetic lethality arises when viable yeast strains carrying deletions of two different genes are crossed to produce inviable, double-mutant progeny (i.e., *gene1Δ* and *gene2Δ* mutant strains are viable separately, but the double-mutant *gene1Δ gene2Δ* strain is inviable). Because they impose strong constraints on the control system, SL gene combinations are exceptionally useful in deducing the network wiring diagram and estimating the rate constants in the mathematical model. On the other hand, if the identification of synthetic lethal combinations of genes is incomplete or inaccurate, then SL ‘identifications’ can wreak havoc on a model by forcing the modeler to make adjustments that are unwarranted. Problems arise because the experimental identification of SL gene combinations is plagued by false-positives and false-negatives and by the fact that some synthetic-lethal interactions are dependent on the genetic background of the parental strain. Hence, for the purpose of modeling cell cycle control in budding yeast, it is crucial to have a reliable, well documented, independently confirmed set of SL gene combinations observed in a uniform genetic background.

We have addressed this problem by reconsidering the identification of SL gene combinations of ‘cell-cycle control’ genes in budding yeast by a disciplined construction of replicate double-mutant strains based on a synthetic gene array (SGA) technology^28^ pioneered by Tong and Boone^29^ and the epistasis miniarray profile (E-MAP) ^28^ workflow described by Schuldiner^30^.

We focused on a set of only 36 cell cycle genes, most of which are functionally well-characterized (Table 1). This list comprises all the non-essential genes included in a recent mathematical model of the yeast cell cycle (herein referred to as the ‘Kraikivski’ model)^19^, as well as genes whose protein products have redundant functions or interact with the proteins represented in the model.

**Table 1.**
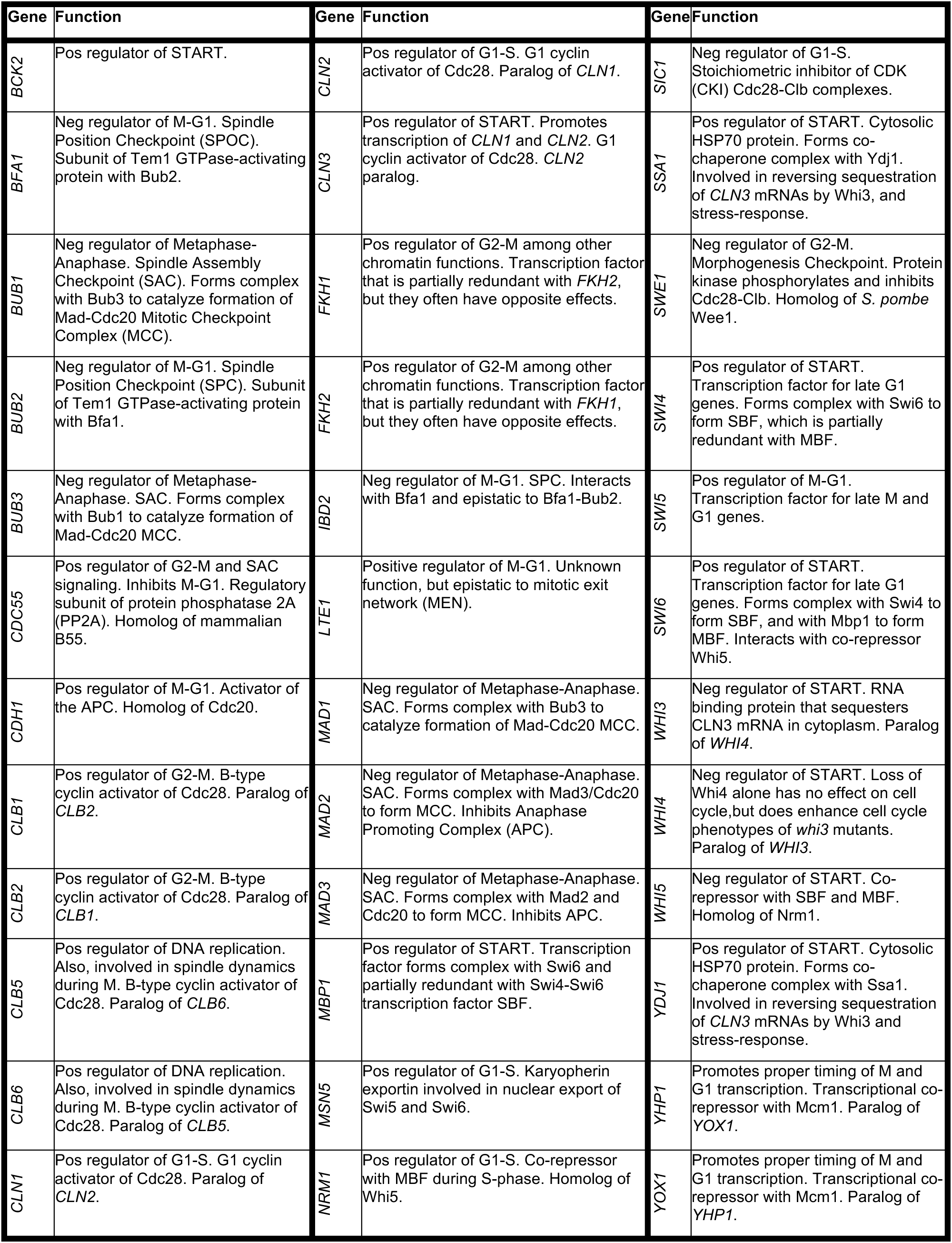
List of 36 cell cycle regulator genes used in the crosses

| Gene | Function | Gene | Function | Gene | Function |
| --- | --- | --- | --- | --- | --- |
| *BCK2* | Pos regulator of START. | *CLN2* | Pos regulator of G1-S. G1 cyclin activator of Cdc28. Paralog of *CLN1*. | *SIC1* | Neg regulator of G1-S. Stoichiometric inhibitor of CDK (CKI) Cdc28-Clb complexes. |
| *BFA1* | Neg regulator of M-G1. Spindle Position Checkpoint (SPOC). Subunit of Tem1 GTPase-activating protein with Bub2. | *CLN3* | Pos regulator of START. Promotes transcription of *CLN1* and *CLN2*. G1 cyclin activator of Cdc28. *CLN2* paralog. | *SSA1* | Pos regulator of START. Cytosolic HSP70 protein. Forms co-chaperone complex with Ydj1. Involved in reversing sequestration of *CLN3* mRNAs by Whi3, and stress-response. |
| *BUB1* | Neg regulator of Metaphase-Anaphase. Spindle Assembly Checkpoint (SAC). Forms complex with Bub3 to catalyze formation of Mad-Cdc20 Mitotic Checkpoint Complex (MCC). | *FKH1* | Pos regulator of G2-M among other chromatin functions. Transcription factor that is partially redundant with *FKH2*, but they often have opposite effects. | *SWE1* | Neg regulator of G2-M. Morphogenesis Checkpoint. Protein kinase phosphorylates and inhibits Cdc28-Clb. Homolog of *S. pombe* Wee1. |
| *BUB2* | Neg regulator of M-G1. Spindle Position Checkpoint (SPC). Subunit of Tem1 GTPase-activating protein with Bfa1. | *FKH2* | Pos regulator of G2-M among other chromatin functions. Transcription factor that is partially redundant with *FKH1*, but they often have opposite effects. | *SWI4* | Pos regulator of START. Transcription factor for late G1 genes. Forms complex with Swi6 to form SBF, which is partially redundant with MBF. |
| *BUB3* | Neg regulator of Metaphase-Anaphase. SAC. Forms complex with Bub1 to catalyze formation of Mad-Cdc20 MCC. | *IBD2* | Neg regulator of M-G1. SPC. Interacts with Bfa1 and epistatic to Bfa1-Bub2. | *SWI5* | Pos regulator of M-G1. Transcription factor for late M and G1 genes. |
| *CDC55* | Pos regulator of G2-M and SAC signaling. Inhibits M-G1. Regulatory subunit of protein phosphatase 2A (PP2A). Homolog of mammalian B55. | *LTE1* | Positive regulator of M-G1. Unknown function, but epistatic to mitotic exit network (MEN). | *SWI6* | Pos regulator of START. Transcription factor for late G1 genes. Forms complex with Swi4 to form SBF, and with Mbp1 to form MBF. Interacts with co-repressor Whi5. |
| *CDH1* | Pos regulator of M-G1. Activator of the APC. Homolog of Cdc20. | *MAD1* | Neg regulator of Metaphase-Anaphase. SAC. Forms complex with Bub3 to catalyze formation of Mad-Cdc20 MCC. | *WHI3* | Neg regulator of START. RNA binding protein that sequesters *CLN3* mRNA in cytoplasm. Paralog of *WHI4*. |
| *CLB1* | Pos regulator of G2-M. B-type cyclin activator of Cdc28. Paralog of *CLB2*. | *MAD2* | Neg regulator of Metaphase-Anaphase. SAC. Forms complex with Mad3/Cdc20 to form MCC. Inhibits Anaphase Promoting Complex (APC). | *WHI4* | Neg regulator of START. Loss of Whi4 alone has no effect on cell cycle, but does enhance cell cycle phenotypes of *whi3* mutants. Paralog of *WHI3*. |
| *CLB2* | Pos regulator of G2-M. B-type cyclin activator of Cdc28. Paralog of *CLB1*. | *MAD3* | Neg regulator of Metaphase-Anaphase. SAC. Forms complex with Mad2 and Cdc20 to form MCC. Inhibits APC. | *WHI5* | Neg regulator of START. Co-repressor with SBF and MBF. Homolog of Nrm1. |
| *CLB5* | Pos regulator of DNA replication. Also, involved in spindle dynamics during M. B-type cyclin activator of Cdc28. Paralog of *CLB6*. | *MBP1* | Pos regulator of START. Transcription factor forms complex with Swi6 and partially redundant with Swi4-Swi6 transcription factor SBF. | *YDJ1* | Pos regulator of START. Cytosolic HSP70 protein. Forms co-chaperone complex with Ssa1. Involved in reversing sequestration of *CLN3* mRNAs by Whi3 and stress-response. |
| *CLB6* | Pos regulator of DNA replication. Also, involved in spindle dynamics during M. B-type cyclin activator of Cdc28. Paralog of *CLB5*. | *MSN5* | Pos regulator of G1-S. Karyopherin exportin involved in nuclear export of Swi5 and Swi6. | *YHP1* | Promotes proper timing of M and G1 transcription. Transcriptional co-repressor with Mcm1. Paralog of *YOX1*. |
| *CLN1* | Pos regulator of G1-S. G1 cyclin activator of Cdc28. Paralog of *CLN2*. | *NRM1* | Pos regulator of G1-S. Co-repressor with MBF during S-phase. Homolog of Whi5. | *YOX1* | Promotes proper timing of M and G1 transcription. Transcriptional co-repressor with Mcm1. Paralog of *YHP1*. |

By comparing the results of our screen with previously published SL interactions listed on The Saccharomyces Genome Database (SGD, https://www.yeastgenome.org/), and further validating via tetrad analysis (TA), we generate lists of ‘high confidence’ and ‘low confidence’ SL interactions. Next, we compare these high-confidence SL interactions with the predictions of our most recent and extensive mathematical model of budding-yeast cell-cycle controls ^19^. We find that, in its present state, the model’s predictions of SL interactions are not very accurate because the predictions were based on parameter values estimated from a collection of SL gene combinations that misidentified some crucial genetic interactions. From our new collection of high-confidence and low-confidence SL gene combinations we re-parametrize the model to get much better agreement with the data. Presumably, this newly parametrized version of the model will give more reliable predictions about the phenotypes of other types of budding yeast mutants as well.

Finally, we phenotype mutants of the ∼600 gene combinations that are not SL under six different media conditions expected to differentially influence cell cycle progression, providing quantitative fitness data that can be used in the future development of more refined and stochastic models of the cell cycle.

## Results

### Identifying synthetic lethal interactions among 630 gene combinations

To assess all possible combinations of 36 cell cycle knock-outs across multiple biological replicates, we generated eight sets of independent parent lines to be used in four crosses. To avoid suppressor mutations – a feature of the commercial yeast haploid gene deletion collections – we generated 110 parent strains by tetrad dissection of commercial heterozygous diploid gene deletion strains (either before or after switching the *kanMX* marker to *natMX*), and we generated 4 parent strains by de novo gene deletion in BY4741 or BY4742. Neither the commercial *SSA1*/*ssa1Δ* strain nor any diploids produced by crosses with any de novo *ssa1Δ* mutant parent were able to sporulate, indicating that two copies of this HSP70 chaperone gene is essential for meiosis. Interestingly this was not the case for the Ssa1 co-chaperone, Ydj1. We also generated SGA haploid selection marker strains by mating and tetrad dissection of the aforementioned strains with the SGA strain developed by the Boone lab^29^ or by de novo gene deletion in that strain (55 and 70 parent strains, respectively).Each set of parent strains carried at least two differently marked deletions in all or most of the 36 genes for each of two different markers. According to the workflow described in Figure 1, single mutant parent strains with opposite markers were crossed and both *MAT***a** and *MATα* double-mutant progeny were selected for using SGA^29^ haploid selection markers, resulting in up to 20 biological replicates for each gene combination. In total, we generated 7,350 mutants in which to analyze the 630 double-mutant combinations.

**Figure 1.**
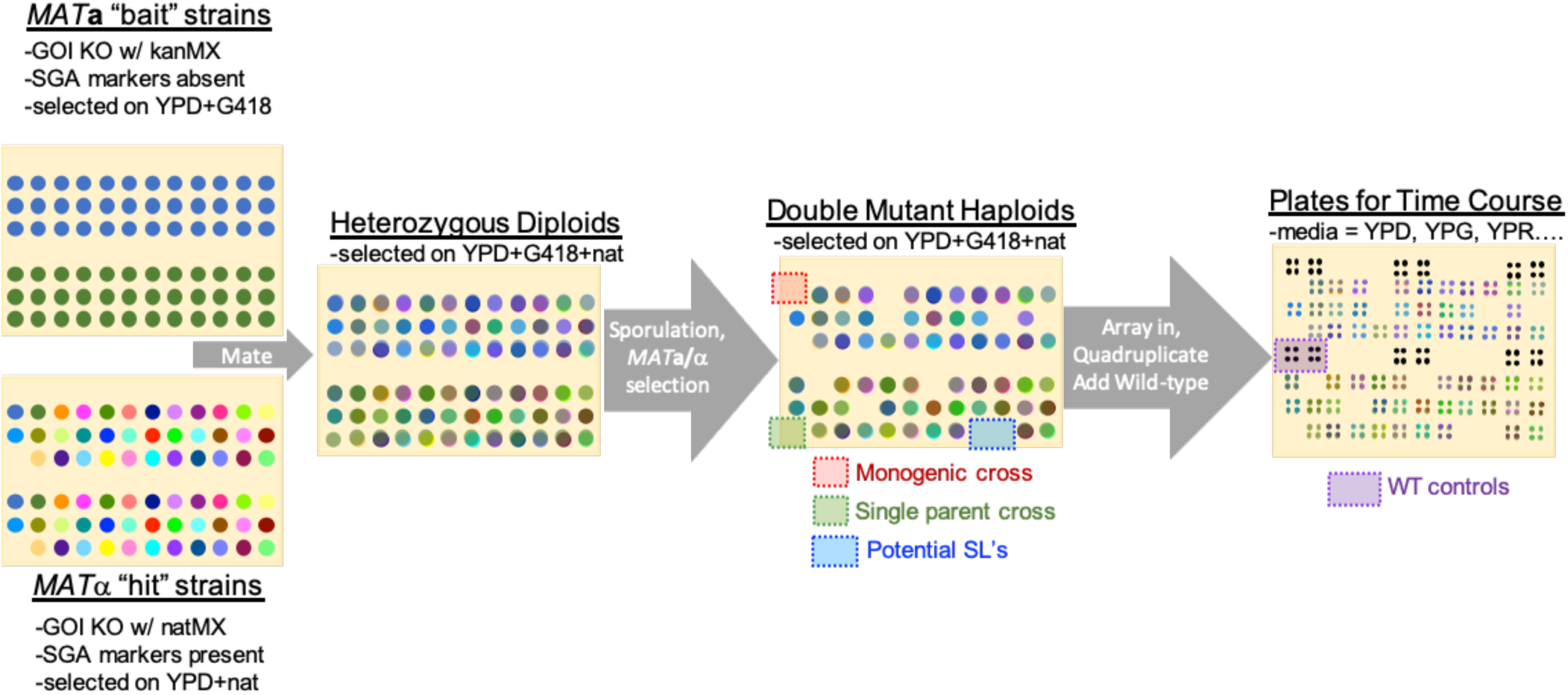
Simplified representation of experimental workflow. Example of a single cross plate where two *MAT***a** “bait” strains in which the gene of interest (GOI) was knocked out (KO) with a kanamycin resistant marker (*kanMX*, also confers resistance to G418) are each crossed to the 36 *MAT***a** “hit” strains in which the gene of interest was knocked out with a nourseothricin-resistant marker (*natMX*). Heterozygous diploids were selected for on media containing both antibiotics, and then sporulated on standard sporulation media. The sporulated colonies were pinned onto a series of specialized SGA media that select for *MAT***a** and *MATα* haploid progeny. Positions on the double mutant haploid plates that would have resulted in “monogenic crosses” (where the same gene of interest was knocked out in both parents) or “single parent crosses” (where one of the parent positions was empty) were monitored for potential false-negatives. The double mutant haploid progeny were used to identify synthetic lethal interactions (Figure 2 and Figure 3) and then pinned in quadruplicate on a fresh YPD plate. WT controls were added, and the resulting master plate was pinned onto six different media types for phenotyping (Figure 4 and Figure 5). Phenotyping plates were imaged every 12 hours to monitor growth rates.

Examining these 7,350 mutants, we first flagged potential synthetic lethal interactions by scoring each cross as ‘growth’ or ‘non-growth’, i.e., each double-mutant haploid colony as ‘present’ or ‘absent’ on double mutant haploid selection plates (Figure 2). The results for all progeny are compiled in Figure 3.

**Figure 2.**
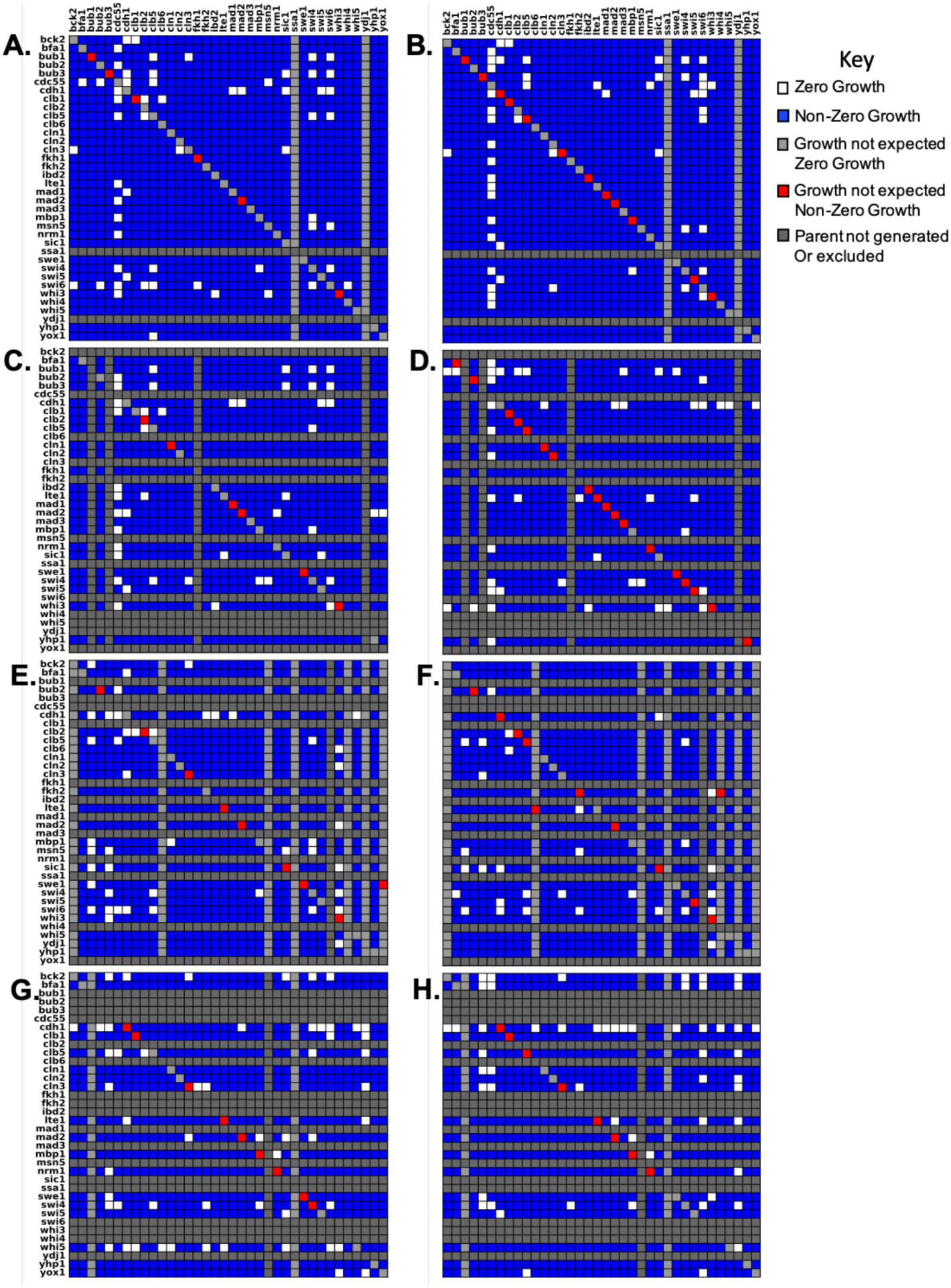
Binary assessment of colony growth for double mutants in all four sets of crosses. Figures on the left were derived from *MAT***a** progeny. Figures on the right were derived from the sister *MATα* progeny. *MAT***a** parents are organized along the x-axis and *MATα* parents are organized along the y-axis alphabetically by the gene that was knocked out. Rows or columns shaded light grey indicate positions on the plate that should have been empty, because the parent was never generated. The diagonal in each heat map indicates a cross between two parents in which the same gene was knocked out. These should not result in growth under selection. Red cells indicate unexpected growth and are an indication of the false negative rate. Rows and columns shaded dark grey indicate parents that were never generated or were excluded from the analysis, because at least one third of the progeny resulting from that parent failed to grow. Duplicates of the same gene/marker combination within the same cross are not shown. Total number of crosses (excluding monogenic) =7350 **A** & **B**) *Cross 1*. **C** & **D**) *Cross 2*. **E** & **F**) *Cross 3.* **G** & **H**) *Cross 4*.

**Figure 3.**
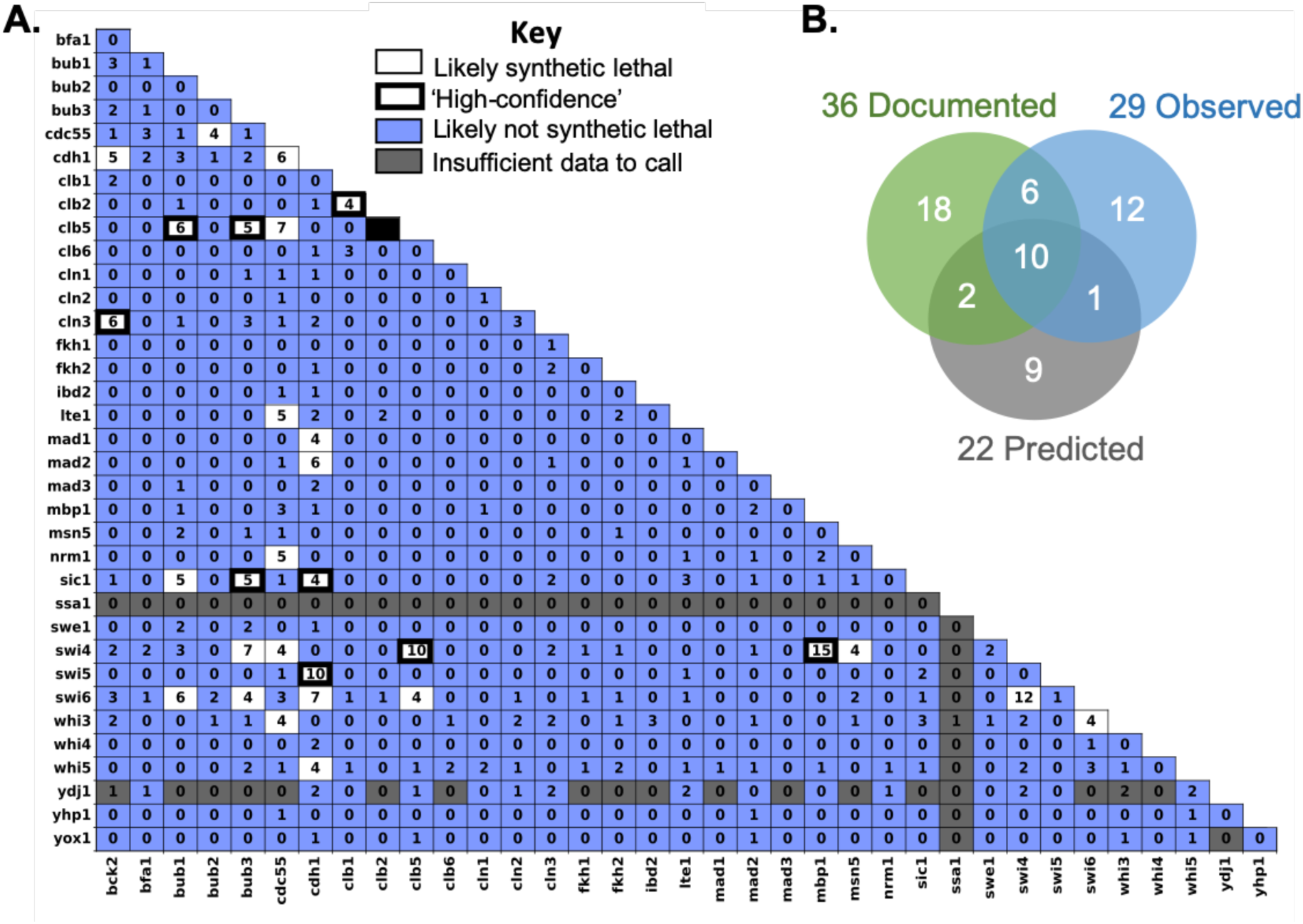
Likely synthetic lethal interactions determined by compiling data from all crosses. **A)** Table documenting how many crosses supported synthetic lethality (no growth of the double mutant progeny). Synthetic lethal interactions that we designate as ‘high-confidence’ in Table 2 are outlined in black **B)** Venn-Diagram comparing observed SL interactions with those that have been previously documented and/or predicted by the Kraikivski model. Note: *clb2Δ clb5Δ* is excluded as these genes are linked.

**Table 2.** Comparison of observed synthetic lethal interactions with those previously documented, those predicted by the Kraikivski model^19^, and our new model. For previously documented SL interactions, it is noted whether they were manually curated or derived from a high-throughput (HTP) screen. The “SL score” highlights the number of crosses in our screen that support synthetic lethality out of the total number of replicates. Double mutant combinations are reported as synthetic lethal (SL), viable (V), or having reduced viability (RV). More than one reported TA result indicates that the TA was repeated for two or more crosses. Results marked with an asterisk are lower confidence due to poor spore viability. Cases where spore viability was too low to determine synthetic lethality are marked MD to indicate a possible meiotic defect. Rows shaded orange mark likely synthetic lethal interactions. Rows shaded blue mark likely viable interactions. Instances where the model does not support the ‘high confidence’ interactions are bolded.

| Gene 1 | Gene 2 | Documented |  | Observed |  |  | Kraikovski Model | New Model |
| --- | --- | --- | --- | --- | --- | --- | --- | --- |
|  |  | Method | Refs | SL | SL Score | TA |  |  |
| BCK2 | CDH1 |  |  | Y | 5/14 | V, V, V | V | V |
| BCK2 | CLN3 | Manual | 33, 77-81 | Y | 6/12 | SL*, SL, RV | SL | SL |
| BCK2 | SWI4 | HTP | 82 |  | 2/16 | V, V, V | V | V |
| BCK2 | SWI6 | Manual | 33 |  | 3/6 | V, V, V | SL | SL |
| BFA1 | BUB3 | HTP | 38 |  | 1/10 | V |  |  |
| BFA1 | LTE1 | Manual | 83 |  | 0/16 | V, V, V |  |  |
| BUB1 | CLB5 | Manual | 31 | Y | 6/8 | SL*, SL* |  |  |
| BUB1 | SIC1 | Manual | 84 | Y | 5/8 | V, SL, MD, V |  |  |
| BUB1 | SWI6 |  |  | Y | 6/10 | RV*, RV* |  |  |
| BUB2 | CDC55 |  |  | Y | 4/8 | V, V | V | V |
| BUB3 | CDH1 | HTP | 62 |  | 2/10 | V |  |  |
| BUB3 | CLB1 | HTP | 62 |  | 0/8 | V |  |  |
| BUB3 | CLB5 | HTP/manual | 31, 62 | Y | 5/10 | SL |  |  |
| BUB3 | SIC1 | HTP | 62 | Y | 5/8 | SL |  |  |
| BUB3 | SWI4 |  |  | Y | 7/12 | V, V |  |  |
| BUB3 | SWI6 |  |  | Y | 4/10 | V, SL |  |  |
| BUB3 | YDJ1 | HTP | 62 |  | 0/2 | V |  |  |
| CDC55 | CDH1 |  |  | Y | 6/10 | SL*, V, MD | V | V |
| CDC55 | CLB5 |  |  | Y | 7/10 | SL, RV | V | SL |
| CDC55 | LTE1 |  |  | Y | 5/10 | SL, RV | V | V |
| CDC55 | MSN5 | Manual | 85 |  | 1/6 | V, V, RV* |  |  |
| CDC55 | NRM1 |  |  | Y | 5/8 | V, V |  |  |
| CDC55 | SWI4 |  |  | Y | 4/12 | V, V, MD | V | V |
| CDC55 | WHI3 |  |  | Y | 4/8 | MD, MD |  |  |
| CDH1 | CLB5 |  |  |  | 0/16 | V | SL | V |
| CDH1 | CLN3 |  |  |  | 2/16 | V | SL | V |
| CDH1 | LTE1 |  |  |  | 2/16 | V, RV, MD | SL | V |
| CDH1 | MAD1 | HTP | 62 | Y | 4/12 | RV, V |  |  |
| CDH1 | MAD2 | HTP | 62 | Y | 6/16 | V, MD, V | SL | SL |
| CDH1 | SIC1 | HTP/manual | 84, 86 | Y | 4/14 | SL | SL | SL |
| CDH1 | SSA1 |  |  |  | 0/2 |  | SL | V |
| CDH1 | SWI4 | HTP | 82 |  | 0/18 | V, V, V | V | V |
| CDH1 | SWI5 | Manual | 87 | Y | 10/16 | SL, RV* | SL | SL |
| CDH1 | SWI6 |  |  | Y | 7/12 | SL, V, MD, MD | SL | SL |
| CDH1 | WHI5 | HTP | 82 | Y | 4/18 | V, RV* | SL | SL |
| CDH1 | YDJ1 |  |  |  | 2/4 | SL, V, RV | SL | V |
| CLB1 | CLB2 | Manual | 88-91 | Y | 4/12 | SL, SL | SL | SL |
| CLB2 | MAD2 | HTP | 62 |  | 0/14 | V | V | V |
| CLB2 | SWI6 |  |  |  | 1/10 | V | SL | V |
| CLB2 | SIC1 | Manual | 92, 93 |  | 0/12 | V, V | V | V |
| CLB5 | SIC1 | Manual | 92, 93 |  | 0/14 | V, V, RV | V | V |
| CLB5 | SWI4 | Manual | 94 | Y | 10/18 | RV*, SL | SL | SL |
| CLB5 | SWI6 | Manual | 94 | Y | 4/12 | V, V, SL | SL | SL |
| CLN3 | SWI4 | HTP, manual | 33, 82 |  | 2/18 | V, V, V, V | SL | V |
| LTE1 | MAD2 |  |  |  | 1/16 | V | SL | V |
| LTE1 | SIC1 | HTP | 84 |  | 3/14 | V*, SL | V | V |
| LTE1 | SWI6 |  |  |  | 1/12 | V | SL | V |
| LTE1 | YDJ1 | HTP | 95 |  | 2/4 | SL, SL, V | V | V |
| MAD1 | SIC1 | HTP | 62 |  | 0/10 | V |  |  |
| MAD2 | SIC1 | HTP | 62 |  | 1/14 | V, V | V | V |
| MAD2 | WHI5 | HTP | 62 |  | 1/18 | V | V | V |
| MAD3 | SIC1 | HTP | 62 |  | 0/10 | V |  |  |
| MBP1 | SWI4 | Manual | 78, 96 | Y | 15/20 | SL | SL | SL |
| MSN5 | SWI4 | Manual | 97, 98 | Y | 4/8 | V |  |  |
| MSN5 | SWI6 | Manual | 97, 98 |  | 2/4 | RV, V |  |  |
| SWI4 | SWI6 | Manual | 33, 99, 100 | Y | 12/14 | RV*, V*, MD | SL | SL |
| SWI4 | YDJ1 |  |  |  | 2/6 | V | SL | V |
| SWI6 | WHI3 |  |  | Y | 4/10 | V, V |  |  |
| Total # SL intractions |  | 36 |  |  | 29 (6 viable by TA) |  | 22 | 13 |

No combination of genes produced the same results in every cross. In fact, the results among biological replicates varied considerably (Figure 2, Table S4). Hence, we set a threshold for defining likely SL interactions. If evidence for synthetic lethality was observed four or more times irrespective of which parent strain the deletions were derived from, we flagged the combination as ‘likely SL’ (Figure 3, Table 2).

A threshold of four was selected, because it ensures that the interaction was seen in at least two of the independent sets of crosses performed. This threshold also provided for the highest level of agreement between our screen and previously published results (discussed in the following section). For our set of 630 combinations, we observed 29 that exhibited synthetic lethality in at least four biological replicates.

### Comparing the results of our screen with previously reported SL interactions

In Table 2 we compare our results to the 36 SL gene combinations documented on SGD for the 36 genes in our data set (excluding several curation errors which are listed in the supplement) and to the predictions of Kraikivski’s published model^19^. There are 58 lines in Table 2, referring to 58 (out of 630 possible) gene combinations for which one or more of the following statements is/are true:

1. the combination is documented as SL on SGD,
2. the combination is observed in our screen as likely SL,
3. the combination has been predicted to be SL by Kraikivski’s model.

A Venn diagram indicating the overlap of these 58 gene combinations is provided in Figure 3B.

Of the 36 gene combinations documented as SL on SGD, 16 were observed as likely SL in our screen (Figure 3B). Meanwhile, 13 of our observed SL gene combinations are not listed on SGD. Hence, the overlap between the previously published SL interactions, and the combinations in our screen that exhibited synthetic lethality in at least four replicates is only ∼50%. Dropping the threshold for likely SL interactions in our screen from four to three would have resulted in the identification of only one additional previously documented SL interaction (*lte1Δ sic1Δ*), while adding 13 SL interactions that are not supported by the literature. Increasing the threshold to five would have excluded an additional 13 SL interactions that have been previously observed.

As a check on these comparisons, we performed tetrad analysis (TA) on at least one cross for all of the combinations listed in Table 2 except for one (*cdh1Δ ssa1Δ*) from which we failed to recover tetrads. Of the 13 SL gene combinations that we observed for the first time in this study, six were not SL by TA (*bck2*Δ *cdh1*Δ, *bub2*Δ *cdc55*Δ, *bub3*Δ *swi4*Δ, *cdc55*Δ *nrm1*Δ, *cdc55*Δ *swi4*Δ and *swi6*Δ *whi3*Δ). The other seven (*bub1*Δ *swi6*Δ, *bub3*Δ *swi6*Δ, *cdc55Δ cdh1Δ, cdc55Δ clb5Δ, cdc55Δ lte1Δ, cd55Δ whi3Δ,* and *cdh1Δ swi6Δ*) exhibited variable results or low spore viability regardless of genotype in at least one of the crosses, complicating the interpretation of the results. Of the 20 ‘documented’ SL interactions that were not observed in our screen, 17/20 tested by TA were viable. The other three (*lte1*Δ *sic1*Δ, *lte1*Δ *ydj1*Δ, *msn5*Δ *swi6*Δ) varied by replicates or exhibited low spore viability overall, and thus remain ambiguous.

In summary, our screen identified 13 new potential SL interactions, but none of these were definitively validated by TA. Our screen failed to validate 20 previously published SL interactions. By TA, we determined that at least 17 of these are likely not SL. Of the 16 double-mutant combinations that were both ‘documented’ SL on SGD and ‘likely’ SL according to our screen, TA confirmed that nine combinations are indeed inviable. The other seven remain ambiguous.

Based on these comparisons, we re-classify the 58 gene combinations in Table 2 as ‘high-confidence synthetic-lethal’ combinations (shaded orange), ‘high-confidence viable’ double-mutants (shaded blue), and ‘uncertain’ (unshaded). Of those that remain uncertain, for five gene combinations which all include *swi4*Δ or *swi6*Δ (*bub3*Δ swi6Δ, *clb5*Δ *swi6*, *msn5*Δ *swi4*Δ, *msn5*Δ *swi6*Δ, *swi4*Δ *swi6*Δ) additional replicates were attempted, but no tetrads were recovered.

Some of the variability observed between replicate tetrad analyses of the same genotype, as well as apparent meiotic defects may be the result of chromosome loss. For instance, Bub1 and Bub3, which are involved in regulating the Spindle Assembly Checkpoint and tension sensing in spindles^31, 32^, exhibited unusual behavior in halo assays indicative of chromosome loss (see additional data).

### Using our screen to refine a previously published model of the cell cycle

In addition to SGD, we compared our ‘likely’ SL interactions with those that were predicted by Kraikivski’s model^19^. Of the 22 predicted SL gene combinations in Table 2, 10 are both documented and confirmed by our screen, two (*bck2Δ swi6Δ* and *cln3Δ swi4Δ*) were documented but not observed by us, and one (*cdh1Δ swi6Δ*) was observed by us but not documented on SGD. Nine predicted SL gene combinations were neither observed by us nor documented on SGD. We tested eight of these by TA and found six to be viable, while two (*cdh1Δ lte1Δ* and *cdh1Δ ydj1Δ*) remain uncertain (Table 2). Five of these ‘orphan’ predictions involve *cdh1Δ*, suggesting an overemphasis of Cdh1 activity in the model. We tested four of these combinations by TA and found that two were viable (*cdh1Δ clb5Δ* and *cdh1Δ cln3Δ*), while two remain uncertain (*cdh1Δ lte1Δ* and *cdh1Δ ydj1Δ*). Six SL gene combinations that were both documented and observed by us were not analyzed in Kraikivski’s model.

In summary, Kraikivski’s model makes 37 predictions (22 SL + 15 V) concerning the genetic interactions listed in Table 2. Of these predictions, 16 are consistent with our ‘high-confidence’ SL/V phenotypes, 7 are inconsistent (bolded in Table 2), and 14 are ambiguous. Hence, the accuracy of the published model is ∼50%, comparable to the agreement between our screen and the literature.

The limited accuracy of the model’s predictions is likely due to the fact that the parameter values in the model were estimated by fitting the model to ‘documented’ SL gene combinations that are themselves unreliable. To correct this problem, we have re-parametrized the model in light of the ‘high confidence’ SL and viable (V) interactions (shaded orange and blue, respectively in Table 2), allowing for some flexibility for the uncertain interactions.

In re-parameterizing the model, we had two intentions: (a) to maximize the number of correctly explained mutant phenotypes in Table 2, and (b) to simulate correctly those mutant strains with well-characterized phenotypes that were previously explained by the model. Guided by these two criteria, we manually adjusted 13 parameter values in the model (see Table S3 in ^19^), as follows:

First, because of the central roles played by SBF, MBF and Cln3 in the START transition of the budding yeast cell cycle, we addressed our new results suggesting a viable phenotype for *swi4Δ cln3Δ* double-mutant cells in opposition to previous reports that *swi4Δ cln3Δ* is a synthetic lethal strain ^33^. To ‘rescue’ *swi4Δ cln3Δ* cells, we significantly increased the activation of MBF (Swi6:Mbp1) by Bck2 (the only activator of MBF in the absence of Cln3), while simultaneously increasing the inactivation of MBF by Clb2 and decreasing slightly the activation of MBF by Cln3, in order to keep the level of MBF activity similar to that of the previous model, thus minimizing the perturbations to all other mutants that were previously explained by the model. Because Ydj1 is a regulator of Cln3 activity, the phenotype of *swi4Δ ydj1Δ* agreed with new data too.

The viability of *swi6Δ clb2Δ* suggests that Swi4 alone (without Swi6) can successfully initiate the START transition, and then the cell cycle can be completed without Clb2 (with Clb1 alone). To correctly simulate this mutant, we had to significantly increase the weight of Swi4 in the transcriptional regulation of the START transition.

We also made adjustments to account for the five mutant strains involving *cdh1Δ* that our original model did not predict correctly. In the model, cell division (upon exit from mitosis) is determined by Clb2 activity dropping below a certain threshold, which is in turn governed by Cdh1 (involved in Clb2 degradation during telophase) and Sic1 (an inhibitor of Clb2-dependent kinase activity as cells return to G1). Hence, the inviability of *cdh1Δ sic1Δ* cells is the crucial mutant defining the cell-cycle exit threshold, and it was correctly predicted by the original model. In this double-mutant, Clb2-dependent kinase activity is down-regulated in anaphase only by Cdc20-dependent degradation of Clb2. (In reality, of course, Clb2 activity depends on many upstream regulators—such as Ydj1, Clb5, Ssa1, Cln3, and Swi6— that affect cell mass at division.) Our new assessment of synthetic lethal interactions allows for better ‘tuning’ of the parameters that govern Clb2 regulation by Cdh1, Cdc20 and Sic1. Additionally, when originally constructing and parametrizing our model, we did not have many *lte1Δ* mutant strains to constrain Lte1-related parameters, so we adjusted parameters to correctly explain *lte1Δ* mutants.

Predictions of the newly parametrized model are given in the last column of Table 2.Our expertise in cell cycle regulation and mutant behavior allowed us to make these parameter adjustments manually; however, computational algorithms for reparameterization may be required if a larger number of novel mutant phenotypes becomes available in the future.

### Inherent limitations of synthetic lethality screens

The SGA process relies on efficient production of double mutant haploid progeny from crosses. Mating defects, low sporulation efficiencies, meiotic defects causing poor spore viability, poor spore germination, or technical problems with the pinning process can prevent the transfer of double mutant cells during haploid selection, resulting in false positives (i.e., poorer growth than there should be; ^29, 34^). Genetic interactions resulting in reduced fitness are also subject to significant selection for genetic mishaps that improve fitness, resulting in false negatives (i.e., better growth than there should be; ^29, 34^). Genetic mishaps resulting in false negatives can include spontaneous mutation to introduce a suppressor mutation ^35^, or disomy. Disomy can result from chromosome nondisjunction during sporulation, or gene conversion resulting in escape of heterozygous diploids from haploid selection ^34, 36^. False negatives can also result from contamination from outside sources or cross-contamination during replica-pinning.

Following the presence or absence of colonies throughout the SGA process, we found that all crosses produced diploids (see Additional Data). Therefore, failure to mate did not produce any false positives. False positives can also result from inefficient pinning or systematic problems with the parents resulting in overall low viability. Parent lines that resulted in fewer than 12/36 viable haploid progeny were excluded from the analysis, but some false positives likely persisted. For instance, seven of the SL gene combinations observed in our screen involved *cdc55Δ*, which was problematic in most genetic contexts due to inconsistent pinning (cells were very dry and clumpy and did not adhere well to pins). By tetrad analysis, we identified six out of 29 SL gene combinations observed in our screen to be definitive false positives.

Our experimental design makes it possible to get rough estimates of false negative rates by monitoring positions on each plate that should have been empty for growth. We designed our screen such that “hit” strains were arrayed the same way (alphabetically by the gene knocked out) for every cross, leaving empty spaces for any parent that was not generated (Figure 1). In this way, some positions in the cross had only one parent crossed to an empty position (‘single parent’ in Figure 1), and some positions had two parents that were mutant for the same allele (‘monogenic cross’ in Figure 1). Neither of these ‘crosses’ should result in colonies during the final round of double-mutant haploid selection, as they will not contain both of the antibiotic resistance markers. Colonies at ‘single parent’ or ‘monogenic cross’ positions are indicative of a false negative event (red cells in Figure 2).

To estimate the contribution of contamination to false negatives, we identified colonies in empty plate positions. All plates were devoid of contaminating colonies in empty positions (Table S3). In positions containing only one parent strain, only 3/570 positions on the haploid progeny plates had any contaminating colonies (Table S3). Therefore, contamination is a negligible contributor to observed false negatives.

To identify false negatives arising from genetic mishaps, we identified colonies produced by crosses between two strains carrying deletions of the same gene. 77/202 monogenic crosses resulted in progeny on the final haploid selection plates (Table S3), indicating a coarsely estimated false negative rate of 38%.

These false negative events occurred more frequently for *MATα* progeny than *MAT***a** progeny (Table S3). This is to be expected, because *MAT***a** progeny can escape selection for *MATα* progeny through gene conversion between *STE3pr-LEU2* and *leu2Δ0*, but gene conversion cannot occur between *STE2pr-SpHIS3* (S. pombe orthologue) and *his3Δ1* to allow *MATα* cells to escape *MAT***a** selection ^29, 34^. If *MAT***a** progeny persist through the *MATα* selection due to gene conversion, they can mate with the neighboring *MATα* progeny producing diploids that are heterozygous for both markers.

Although few SGA or E-MAP studies report them, it is well-established that these screens have high, but variable, false negative and false positive rates from 17% to 70% ^30, 36–39^ and 5% to 90% ^38, 40–43^, respectively. The false positive and negative rates observed in our study are thus in the normal range for large genetic screens.

### Quantifying fitness and genetic interactions across six media types

As the most extreme genetic interaction, synthetic lethality has a powerful influence on models of cell-cycle regulating genes. However, due to the limitations of synthetic lethality screens more accurate models call for more nuanced phenotypic markers.

To identify interactions between the 36 genes that do not result in synthetic lethality, we monitored the growth rate of the viable double mutants over a time course. Each mutant was assigned a fitness score according to how the growth rate compared with wild-type controls on the same plate. Using this approach, we identified ∼100 gene combinations that were not SL but had fitness scores more than six standard deviations below wild-type under normal growth conditions (Figure 4).

**Figure 4.**
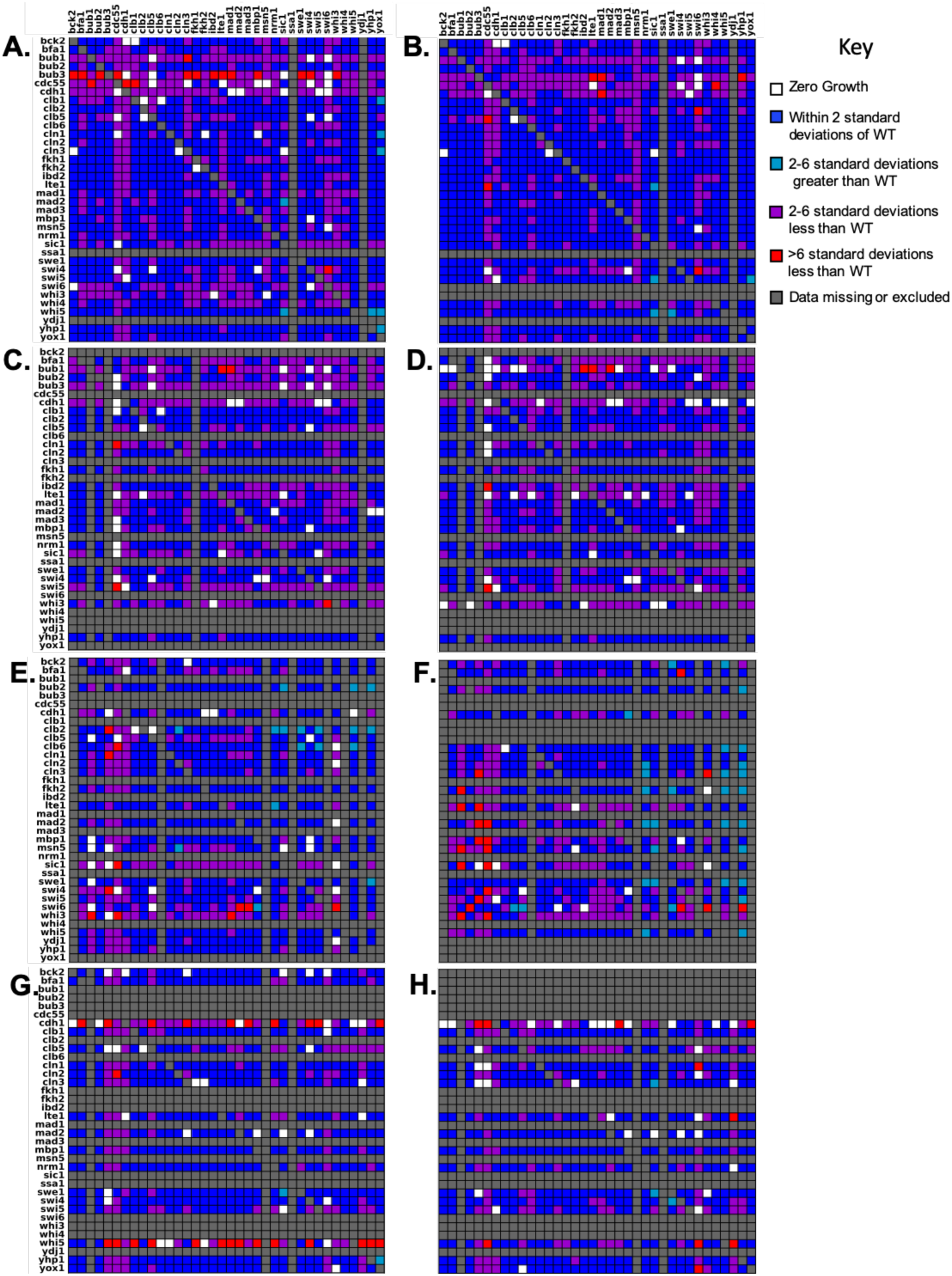
Comparison of fitness scores for double mutants in all four sets of crosses on YPD media. Figures on the left were derived from *MAT***a** progeny. Figures on the right were derived from the sister *MATα* progeny. *MAT***a** parents are organized along the x-axis and *MATα* parents are organized along the y-axis alphabetically by the gene that was knocked out. Rows and columns shaded dark grey indicate crosses that were not performed or were excluded from the analysis. **A** & **B**) Cross 1 **C** & **D**) Cross 2. **E** & **F**) Cross 3 **G** & **H**) Cross 4. Fitness heatmaps for the remaining 5 media types are available is Figures S3-S7. Duplicates of the same gene/marker combination within the same cross are not shown.

More importantly, by comparing fitness scores of the double mutant progeny with those of their single mutant parents, we calculated genetic interaction (GI) scores for all viable mutants. GI scores^44–46^ are a function of the parent and progeny fitness and illustrate the direction (positive or negative) and the magnitude of the interaction for each of the ∼600 viable gene combinations. Non-zero GI scores indicate a possible epistatic relationship. Negative GI scores suggest that the genes involved may have redundant functions, while positive GI scores indicate that one mutation may have a rescuing effect over the other.

As with synthetic lethality, we observed a considerable amount of variability in fitness scores and genetic interaction scores for mutants of the same genotype in different crosses (biological replicates, Figure 4). To identify trends within the variability, GI scores for a given genotype were sorted into different bins, and the bin that contained the largest number of biological replicates was used to determine a consensus GI score which is represented in Figure 5 and Table 3. From the distribution of overall GI scores for a given media, we flagged those with a consensus score at the extreme positive and negative ends. Those gene combinations with consensus GI scores in the top or bottom 5% of all GI scores are reported in Table 3.

**Figure 5.**
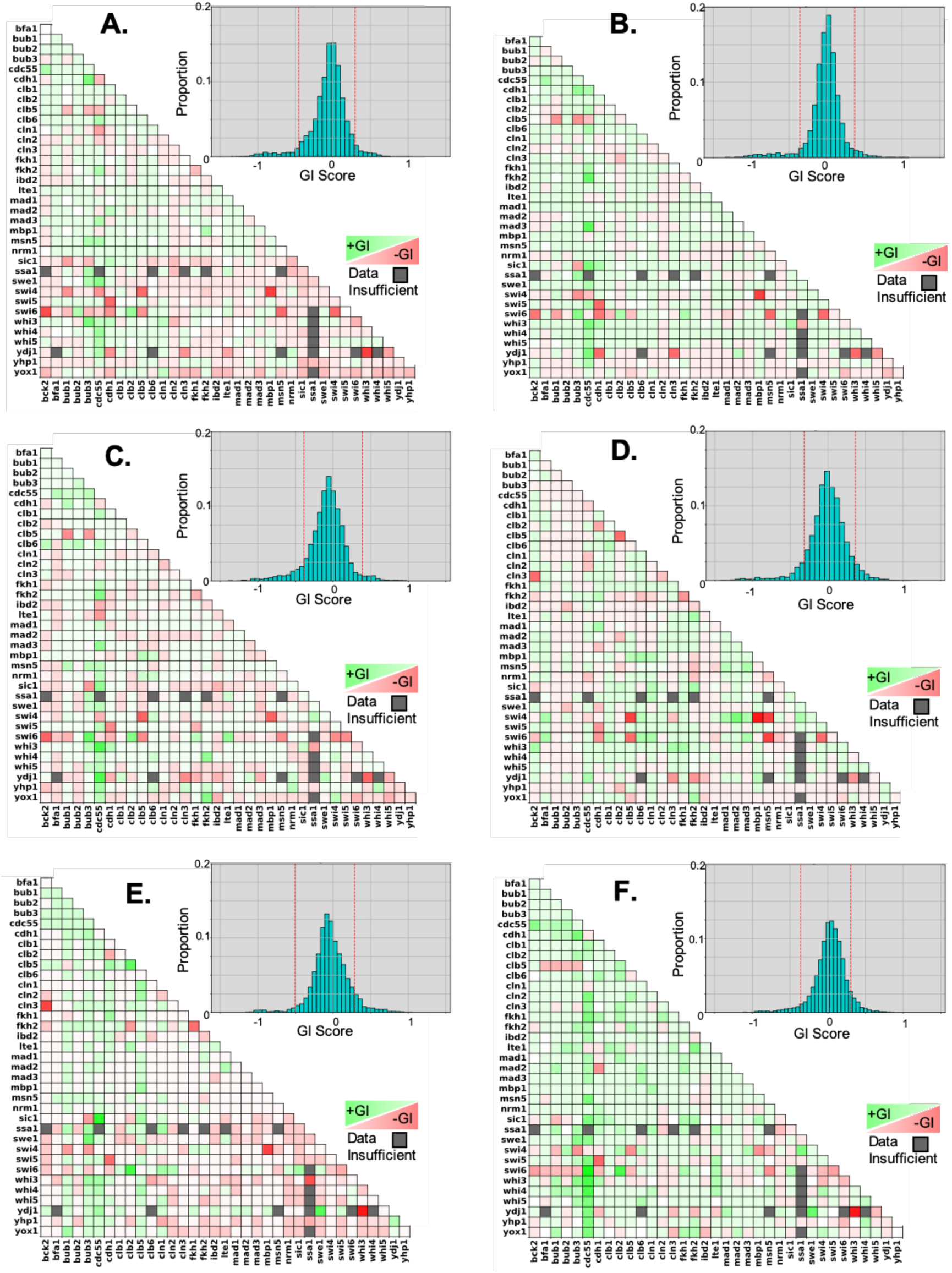
Genetic interaction scores on each media type determined by compiling data from all crosses. Heat maps show the distribution of binned genetic interaction (GI) scores for each mutant combination. Brighter green and darker red squares correspond to higher positive and lower negative GI scores, respectively. Grey squares denote gene combinations for which three or fewer crosses were generated. Histograms show the overall distribution of GI scores for each media type. The dotted red lines distinguish the lower and upper 5% of interactions. **A**: YPD; **B**: YPD + raffinose; **C**: YPD + galactose, **D**: YPD + benomyl; **E**: YPD + camptothecin, **F**: YPD + hydroxyurea

**Table 3.** Genetic interaction score outliers for each media type. Consensus genetic interaction (GI) scores are reported for each gene combination that had a score in the top or bottom 5% of overall GI scores for one or more media types. The number reported reflects the midpoint of the bin occupied by the largest number of biological replicates. Red and green shading highlights negative and positive scores, respectively. The top 2.5% and 5% are shaded light and dark green, respectively. The bottom 2.5% and 5% are shaded light and dark red, respectively. Interactions which we determined to be very likely synthetic lethal are shown in bold. “ND” indicates GI scores that were not determined.

|  |  | YPD | YPR | YPG | YPD+Ben | YPD+CPT | YPD-HU |
| --- | --- | --- | --- | --- | --- | --- | --- |
| BCK2 | CDC55 | 0.327 | 0.126 | 0.040 | 0.163 | 0.175 | 0.401 |
| **BCK2** | **CLN3** | -0.113 | 0.126 | 0.088 | -0.797 | -0.834 | 0.084 |
| BCK2 | SWI6 | -0.918 | -0.703 | -0.824 | -0.317 | -0.229 | -0.397 |
| BCK2 | WHI3 | 0.107 | 0.126 | 0.040 | 0.403 | -0.027 | 0.084 |
| BFA1 | CDC55 | 0.107 | 0.347 | 0.256 | -0.077 | 0.175 | 0.202 |
| BFA1 | SWI4 | -0.113 | -0.095 | -0.176 | 0.403 | -0.027 | 0.084 |
| BFA1 | SWI6 | -0.113 | 0.126 | -0.295 | 0.403 | -0.027 | -0.279 |
| **BUB1** | **CLB5** | -0.332 | -0.537 | -0.608 | -0.077 | -0.138 | -0.320 |
| BUB1 | SIC1 | -0.475 | -0.095 | -0.057 | -0.077 | -0.027 | -0.279 |
| BUB1 | SWI6 | -0.552 | -0.095 | -0.392 | -0.077 | -0.431 | -0.279 |
| BUB1 | SWI6 | -0.332 | -0.460 | 0.040 | -0.077 | 0.377 | 0.265 |
| BUB1 | YOX1 | 0.107 | -0.095 | 0.040 | 0.055 | 0.377 | -0.279 |
| BUB2 | SWI6 | 0.327 | -0.095 | 0.180 | -0.077 | 0.084 | 0.472 |
| BUB3 | CDH1 | 0.547 | 0.445 | -0.532 | -0.077 | 0.306 | 0.472 |
| **BUB3** | **CLB5** | -0.332 | -0.537 | -0.532 | -0.077 | -0.027 | -0.279 |
| BUB3 | MSN5 | 0.327 | 0.126 | 0.256 | -0.077 | 0.175 | 0.084 |
| **BUB3** | **SIC1** | -0.332 | -0.442 | -0.532 | -0.077 | -0.431 | 0.027 |
| BUB3 | SSA1 | 0.341 | 0.3914 | 0.256 | 0.143 | 0.389 | 0.276 |
| BUB3 | SWE1 | 0.327 | 0.126 | 0.180 | -0.077 | 0.579 | 0.401 |
| BUB3 | SWI4 | -0.113 | -0.551 | 0.040 | -0.077 | -0.431 | -0.461 |
| BUB3 | SWI6 | -0.332 | 0.513 | 0.616 | -0.077 | -0.229 | -0.279 |
| BUB3 | WHI3 | 0.547 | 0.126 | 0.256 | 0.055 | 0.377 | 0.401 |
| CDC55 | CDH1 | -0.332 | 0.347 | -0.176 | -0.209 | -0.027 | -0.098 |
| CDC55 | CLB6 | 0.327 | 0.347 | 0.256 | 0.163 | 0.175 | 0.084 |
| CDC55 | CLN1 | -0.475 | 0.126 | 0.088 | -0.317 | -0.027 | -0.098 |
| CDC55 | CLN2 | -0.233 | 0.027 | -0.176 | -0.317 | -0.027 | 0.447 |
| CDC55 | FKH1 | 0.107 | 0.347 | 0.256 | -0.077 | -0.027 | 0.447 |
| CDC55 | FKH2 | 0.327 | 0.665 | 0.472 | 0.163 | 0.175 | 0.276 |
| CDC55 | LTE1 | -0.113 | -0.095 | -0.532 | -0.077 | -0.027 | -0.098 |
| CDC55 | MAD3 | 0.327 | 0.716 | 0.418 | -0.077 | -0.027 | 0.265 |
| CDC55 | MSN5 | 0.327 | 0.108 | 0.256 | 0.163 | -0.027 | 0.447 |
| CDC55 | SIC1 | 0.107 | 0.568 | 0.418 | -0.077 | 0.898 | 0.447 |
| CDC55 | SWE1 | 0.492 | 0.347 | 0.256 | 0.163 | 0.175 | 0.401 |
| CDC55 | SWI4 | -0.552 | -0.316 | -0.392 | -0.209 | -0.027 | -0.098 |
| CDC55 | SWI5 | 0.107 | 0.347 | 0.256 | 0.163 | 0.175 | 0.447 |
| CDC55 | SWI6 | -0.233 | -0.095 | -0.176 | 0.163 | -0.027 | 0.810 |
| CDC55 | WHI3 | 0.250 | 0.568 | 0.688 | 0.163 | 0.377 | 0.628 |
| CDC55 | WHI4 | 0.425 | 0.347 | 0.256 | 0.163 | 0.306 | 0.628 |
| CDC55 | WHI5 | 0.327 | 0.347 | 0.256 | 0.163 | 0.175 | 0.447 |
| CDC55 | YDJ1 | 0.327 | 0.568 | 0.688 | 0.403 | 0.377 | 0.447 |
| CDC55 | YHP1 | 0.156 | 0.347 | 0.352 | -0.077 | 0.175 | 0.447 |
| CDC55 | YOX1 | -0.113 | 0.126 | 0.256 | 0.163 | 0.175 | 0.447 |
| CDH1 | CLB2 | -0.113 | -0.095 | 0.040 | -0.473 | -0.344 | 0.084 |
| CDH1 | CLB5 | 0.107 | 0.126 | 0.040 | 0.163 | 0.377 | 0.084 |
| CDH1 | FKH1 | 0.008 | 0.027 | 0.040 | 0.163 | 0.377 | 0.265 |
| CDH1 | MAD1 | 0.008 | 0.057 | -0.176 | 0.403 | 0.084 | -0.098 |
| CDH1 | MAD2 | 0.250 | 0.126 | 0.040 | 0.055 | -0.027 | -0.461 |
| **CDH1** | **SIC1** | 0.250 | 0.347 | 0.040 | -0.077 | -0.027 | 0.265 |
| **CDH1** | **SWI5** | -0.772 | -0.758 | -0.608 | -0.557 | -0.6325 | -0.642 |
| CDH1 | SWI6 | -0.552 | -0.703 | -0.392 | -0.557 | -0.229 | -0.279 |
| CDH1 | WHI3 | 0.250 | 0.347 | 0.256 | 0.163 | 0.175 | 0.265 |
| CDH1 | YDJ1 | -0.552 | -0.758 | -0.392 | -0.077 | -0.027 | -0.279 |
| CLB2 | CLB5 | 0.008 | -0.095 | -0.176 | -0.797 | 0.667 | 0.265 |
| CLB2 | FKH2 | -0.113 | -0.095 | -0.176 | -0.209 | -0.431 | -0.098 |
| CLB2 | SWI6 | 0.425 | 0.347 | 0.472 | 0.163 | 0.780 | 0.628 |
| CLB5 | CLN1 | 0.107 | 0.126 | -0.057 | 0.163 | 0.377 | 0.084 |
| CLB5 | CLN2 | 0.107 | 0.126 | 0.040 | 0.163 | 0.377 | 0.265 |
| CLB5 | MBP1 | 0.107 | 0.126 | 0.256 | 0.319 | 0.377 | 0.124 |
| CLB5 | SIC1 | -0.113 | 0.126 | -0.176 | 0.163 | 0.377 | 0.084 |
| **CLB5** | **SWI4** | -0.772 | 0.126 | -0.824 | -1.037 | -0.229 | -0.461 |
| CLB5 | SWI6 | -0.552 | -0.316 | -0.608 | -0.797 | -0.027 | -0.279 |
| CLB6 | MBP1 | 0.107 | 0.126 | 0.180 | 0.403 | -0.027 | 0.084 |
| CLB6 | SWI6 | 0.107 | 0.126 | 0.040 | 0.403 | 0.084 | 0.084 |
| CLB6 | WHI3 | 0.107 | 0.126 | 0.040 | 0.163 | -0.360 | -0.098 |
| CLN1 | SSA1 | 0.107 | 0.126 | -0.057 | 0.319 | 0.377 | 0.265 |
| CLN1 | SWI6 | 0.107 | 0.126 | 0.040 | 0.403 | 0.377 | 0.084 |
| CLN1 | YHP1 | -0.113 | -0.095 | -0.176 | 0.163 | -0.360 | 0.084 |
| CLN3 | WHI3 | -0.113 | 0.126 | 0.040 | 0.403 | -0.027 | -0.098 |
| CLN3 | YDJ1 | -0.552 | -0.758 | -0.608 | -0.557 | -0.027 | -0.279 |
| FKH1 | FKH2 | -0.332 | -0.095 | -0.392 | -0.557 | -0.6325 | 0.084 |
| FKH1 | SWI6 | 0.107 | 0.347 | 0.256 | 0.163 | -0.138 | 0.265 |
| FKH1 | WHI3 | 0.107 | 0.126 | 0.040 | 0.403 | -0.027 | 0.084 |
| FKH2 | LTE1 | -0.113 | 0.126 | 0.040 | 0.403 | 0.084 | 0.401 |
| FKH2 | MBP1 | 0.327 | 0.126 | 0.040 | -0.209 | 0.084 | 0.002 |
| FKH2 | SIC1 | -0.233 | 0.126 | -0.176 | 0.403 | -0.027 | 0.265 |
| FKH2 | WHI4 | -0.113 | 0.126 | 0.418 | 0.055 | 0.084 | 0.084 |
| FKH2 | YDJ1 | -0.113 | -0.095 | -0.176 | -0.557 | -0.027 | -0.279 |
| FKH2 | YOX1 | 0.107 | -0.095 | 0.472 | 0.403 | -0.166 | 0.265 |
| LTE1 | SWI6 | 0.156 | -0.095 | 0.418 | -0.209 | -0.027 | 0.265 |
| LTE1 | YDJ1 | -0.458 | -0.095 | -0.176 | -0.077 | -0.027 | -0.223 |
| MAD1 | SWI4 | -0.113 | 0.126 | 0.040 | 0.403 | -0.027 | 0.084 |
| MAD2 | SWI4 | 0.107 | 0.126 | -0.057 | 0.643 | -0.027 | 0.084 |
| MAD3 | MSN5 | -0.113 | 0.027 | 0.040 | 0.403 | -0.027 | 0.084 |
| MAD3 | SWI4 | -0.113 | 0.027 | -0.176 | 0.403 | -0.027 | 0.265 |
| MAD3 | SWI6 | 0.107 | 0.126 | -0.057 | 0.403 | 0.084 | -0.098 |
| **MBP1** | **SWI4** | -0.991 | -0.979 | -0.824 | -1.277 | -0.834 | -0.461 |
| MSN5 | SWI4 | -0.113 | -0.095 | -0.176 | -1.037 | -0.229 | -0.279 |
| MSN5 | SWI6 | -0.772 | -0.758 | -0.608 | -1.037 | -0.229 | -0.461 |
| SIC1 | SWI6 | -0.113 | 0.126 | 0.040 | -0.077 | 0.377 | 0.256 |
| SIC1 | WHI3 | 0.107 | 0.347 | 0.180 | -0.077 | -0.027 | 0.084 |
| SSA1 | SWI4 | -0.113 | 0.126 | 0.256 | 0.403 | -0.037 | 0.447 |
| SSA1 | WHI3 | ND | -0.399 | -0.473 | ND | -0.767 | ND |
| SSA1 | YHP1 | -0.332 | -0.095 | -0.176 | -0.077 | -0.360 | -0.098 |
| SWE1 | YDJ1 | 0.008 | 0.126 | 0.256 | 0.163 | 0.713 | 0.526 |
| SWI4 | SWI6 | -0.772 | -0.758 | -0.608 | -0.797 | -0.229 | -0.279 |
| SWI5 | SWI6 | 0.008 | -0.095 | -0.608 | -0.077 | -0.360 | -0.397 |
| WHI3 | YDJ1 | -0.991 | -0.758 | -0.824 | -0.557 | -1.0361 | -1.0053 |
| WHI5 | YDJ1 | -0.475 | -0.537 | -0.392 | -0.077 | -0.027 | -0.279 |

To further identify genetic interactions among our set of cell cycle regulator genes that may not be apparent under standard growth conditions, we also calculated fitness and genetic interaction scores for all double mutant progeny and single mutant parents in the presence of two different carbon sources and in the presence of three checkpoint activating drugs.

YPDextrose served as a control (mass doubling time ∼100 min). Mass doubling times are longer on YPGalactose (∼150 min) and even longer on YPRaffinose (∼200 min) ^47^. Slower growth rates can enable positive regulators to build up such that mutants which would normally grow very slowly due to the stochasticity of cell cycle transitions can exhibit some level of rescue on YPG or YPR ^23, 48^.

The distribution of GI scores that we observed was comparable for YPD and YPG, but the GI scores occupied a narrower range for mutants grown on YPR (Figure 5), suggesting that the very slow growth rate provided by YPR might allow the growth of mutants that have more extreme phenotypes on YPD to normalize on YPR.

The drugs Benomyl (Ben), camptothecin (CPT), and hydroxyurea (HU) activate checkpoints ^49–59^. Mutants defective in these checkpoints will rush through the cell cycle and accumulate genetic/chromosome defects leading to slower growth due to decreased viability. We expect known checkpoint mutants to exhibit reduced fitness under these conditions, but interactions with other cell cycle regulators (including other checkpoint genes) can enhance or suppress the checkpoint defects ^31, 60–66^.

In several cases, gene combinations that had a GI score within the normal distribution on YPD, showed a much more extreme GI on one or more of the other five media types (Table 3). For instance, the GI score for *fkh1Δ fkh2Δ* on YPD was negative, but not remarkably so. However, on YPD+Ben and YPD+CPT, the consensus GI scores for *fkh1Δ fkh2Δ* were in the lower 5% and 2.5%, respectively. Fkh1 and Fkh2 both promote the transition from G2 to M, so the double mutant is likely to cause stalling at G2. Ben prevents spindle assembly while activating the spindle assembly checkpoint, so that cells move forward to M phase despite not properly forming a mitotic spindle. CPT causes DNA damage during M phase. So, cells that make it to M phase in an *fkh1Δ fkh2Δ* mutant would likely arrest in the presence of Ben or CPT, thus exacerbating the mutant phenotype.

Relative GI scores for a family of gene combinations also reflect the role of those genes within the cell-cycle regulatory network. For instance, Bub1 and Bub3 function along with Mad1, Mad2, and Mad3 to arrest cells in metaphase in response to defective attachments of kinetochores to spindle microtubules – a mechanism called the Spindle Assembly Checkpoint (SAC)^31, 32^. However, Bub1 and Bub3 also have a role in tension sensing in spindles independent of their role in the SAC^31, 32^. This can be seen in the observation that *bub1/3* mutants have lower GI scores in benomyl than mad1-3 mutants (Table 3).

Interestingly, several other mutants did not show reduced fitness in benomyl but did display a chromosome loss phenotype (Table 3, Additional Data). These mutants were also synthetic lethal or synthetic sick with *bub1/3* mutants. Clb5 is one such mutant and has previously been predicted to have a role in tension sensing^31^, which the genetic interaction suggests works independently of Bub1/Bub3. Interestingly, although Sic1 works to inhibit CDK-Clb^67–69^, including Clb5, the *sic1Δ* phenotypes were similar to those of *clb5* mutants. Since Sic1 is important for suppressing CDK/Clb activity and is activated by the mitotic exit network (MEN), we hypothesize that elevated CDK/Clb may prolong anaphase resulting in spindle positioning defects, or defects in SAC silencing.

Although slow growth of *swi6Δ* mutants made it difficult to assess halos, like *clb5Δ* and *sic1Δ* mutants, they also appeared to increase chromosome loss. However, unlike Clb5 and Sic1, Swi6 has no direct role in mitosis. Nevertheless, reduced viability in *bub1/3 swi6* double mutants suggests some interaction. We propose that reduced activity of the MBF and SBF at START perturbs expression of proteins important for spindle function or chromosome cohesion, exacerbating the chromosome segregation defects of the *bub1/3* mutants.

It is important to note that not all of the gene combinations that we identified as ‘high-confidence’ synthetic lethal had remarkably negative genetic interaction scores in our screen. There are two plausible explanations for this discrepancy. First, the use of in-plate wild-type controls prohibited the use of antibiotics in the phenotyping screen, so false negatives (growth where growth is not expected) due to contamination are more likely. Second, for gene combinations that are truly synthetic lethal, any living colonies are necessarily the result of false negatives due to genetic mishaps. These gene combinations are thus more prone to result in outliers with higher than expected genetic interaction scores and should be interpreted with caution.

## Discussion

The selective pressure applied by synthetic lethal screens leads to genetic mishaps that enable mutants that would otherwise be lethal to escape^29, 34^; conversely, the low fitness of many of the single mutant parents used in such screens can cause interactions that are not lethal to emerge. These false negative and false positive events lead to very high levels of variability (see Table S5 for an example). We accounted for this variability by probing a relatively small number of genes with an unprecedented number of biological replicates. While E-MAP screens generally incorporate four biological replicates^24^, and SGA screens rely on technical replicates alone^29^, most of the genetic interactions tested in this study included between eight and 16 independent biological replicates (Table S4). We also compared our results with previous publications and resolved discrepancies via tetrad analysis in order to generate a list of ‘high confidence’ synthetic lethal interactions which informed a new iteration of a previously published cell cycle model.

Variability in synthetic lethal screens is a major challenge for modelers. The ∼100 tetrad analyses performed in this study demonstrate an unexpectedly high level of variation even among low-throughput, manual experiments. For this reason, synthetic lethality may not be the best marker for parameterizing models. Additionally, models based on synthetic lethality are inherently deterministic; yet, it’s well-known that many of the processes governing progression through the cell cycle are stochastically regulated. Modeling stochasticity will require a more granular dataset that provides quantitative phenotypes based on parameters such as growth rate, rather than deterministic phenotypes such as lethality or checkpoint arrest.

The results presented here demonstrate that quantitative cell phenotyping can be readily performed in a high-throughput workflow. By comparing colony sizes over time, we generated a quantitative picture of growth rates for over 7,000 mutants. This more sensitive approach enabled us to identify interesting genetic interactions with less extreme phenotypes than synthetic lethality (ie. *whi3Δ ydj1Δ*) and gene combinations that provided a rescue effect (ie. *bub3Δ cdh1Δ*). We also show that our workflow can be expanded to include different test conditions. By quantitatively phenotyping our mutants on six different media types, we demonstrate that our approach is sensitive enough to capture environmental variability. Data for the ∼44,000 gene by media combinations is available through the supplement and can be used to develop more elaborate models of cell cycle regulatory control.

In conclusion, our approach is readily scalable and could generate additional, multi-gene data sets to motivate the development of better, more stochastic models of cell-cycle control and, indeed, other aspects of the physiology of budding yeast cells.

## Methods

### Experimental Workflow

To generate the double mutants, we used a modified epistasis miniarray profile (E-MAP) workflow ^30^. The E-MAP workflow is a modification of the synthetic genetic array (SGA) protocol ^30^. In a typical SGA screen, a single query strain is crossed to all viable deletion strains (over 4,000) ^29, 40^. The query strain includes a set of reporter genes that allow selection of haploid progeny of one mating type or another. E-MAP screens use the same series of selection conditions, but generally involve a few hundred deletion strains crossed to produce every possible combination of double-gene deletions ^30^.

Our experimental design most closely follows the E-MAP approach but with a few significant differences. First, we focused on a set of only 36 cell cycle genes. Second, we used eight sets of parent strains in four sets of crosses, increasing the number of biological replicates to eight from four in a standard E-MAP or one in a standard SGA (which use technical replicates; ^29, 30^):

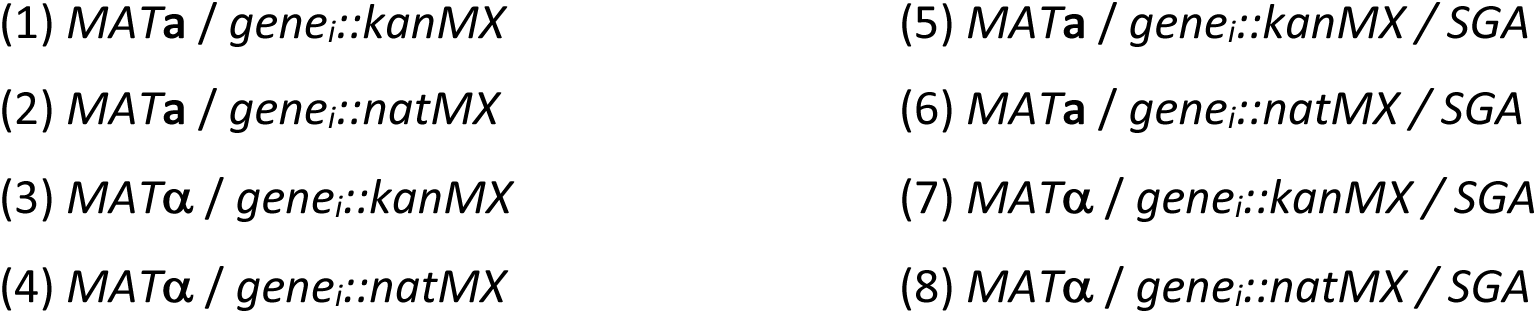

*gene_i_::kanMX* refers to *gene_i_* knocked-out with a kanamycin-resistance marker, *gene_i_::natMX* refers to *gene_i_*knocked-out with a nourseothricin-resistance marker, and ‘*SGA*’ refers to the haploid-selection markers *can1Δ::STE2pr-Sphis5* and *lyp1Δ::STE3pr-LEU2* used in SGA screens. Details for how each of the parent strains were generated can be found in the online supplement. These parent strains were confirmed by PCR and used in four sets of crosses:

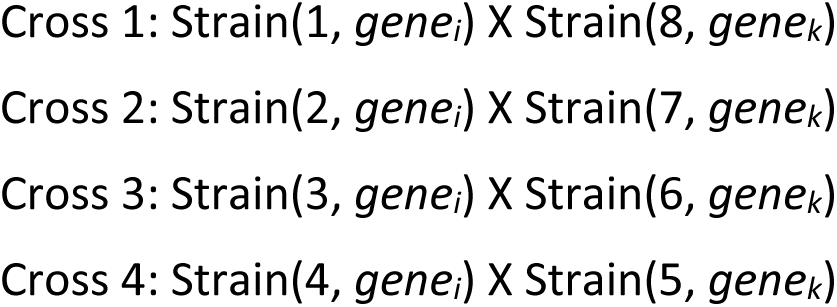

From these crosses, we selected double-mutant progeny of both mating types, further increasing the biological replicates to 16. (SGA and E-MAP screens select only *MAT*a progeny ^29, 30^). All media were standard recipes for SGA^29^ (see Supplemental Materials). Mating type of the double mutant progeny and the single mutant parents was confirmed via Halo assays^70^ (See online supplement). With this protocol, we generate (in principle) 16 biological replicates of each double-mutant, *gene_i_Δ gene_k_Δ*. In certain cases, two parents of the same genotype were generated independently, such that the total number of biological replicates is up to 20 for some double mutant combinations.

We measured colony growth rates on three different growth media (YPD, YPG, and YPR) and in the presence of three different checkpoint activating drugs: Ben, CPT, and HU. Ben disrupts attachment of kinetochores to the mitotic spindle and activates the spindle assembly checkpoint (SAC; dependent on Bub1,3 and Mad1-3) ^49, 71, 72^. CPT inhibits topoisomerase resulting in DNA entanglements and double strand breaks upon chromosome segregation ^73, 74^. HU inhibits ribonucleotide reductase ^75^, which leads to replication fork stalling ^76^.

We first made template plates by first replica pinning the haploid progeny (which were in 96 array) onto different positions on the same YPD+G418(600ug/ml)/nat(150ug/ml) source plate four times to produce quadruplicates of each strain using a Rotor HDA (Singer Instruments, Somerset, UK). Rows A, B, I and J were left empty for in-plate wild-type controls colonies. At the same time, we set up YPD plates with the wild-type parent strains BY4741 and BY4742 arrayed at 384 density, occupying positions in rows A, B, I and J. We incubated both sets of plates at 30°C for two days.

We then replica pinned the wild-type controls onto new YPD plates (using a new source plate whenever the colonies began to look depleted). After visual inspection of the plates to ensure even transfer of the wild-type controls, we replica pinned the set of double mutant colonies to the templates. Plates were imaged after 12, 24, 36, 48, and 60 hours of growth at 30°C.

We imaged all diploid selection plates, final haploid progeny selection plates, halo assay plates, and phenotyping plates using the Phenobooth (Singer Instruments, Somerset, UK) imaging platform and software. To maintain consistency, all images were collected in the same order at the same resolution and camera settings, and were batch processed to crop the image, perform background subtraction and colony identification whenever possible. We then exported the raw colony size data for analysis.

### Data Analysis

Analysis of the data is discussed extensively in the supplement. Briefly, plate-to-plate variation was accounted for by normalizing colony size using in-plate wild-type controls. Edge-effects were accounted for by adjusting the growth rates such that the mean growth rates of edge-adjacent colonies and internal colonies were comparable (Table S6, Figure S1, and Figure S2). Jack-knife filtering was used in a small number of cases to remove colonies that behaved as outliers within quadruplicates (four technical replicates).

Growth rates, fitness scores, and GI scores^44–46^ were calculated using a linear model for growth rate according to the following equations:

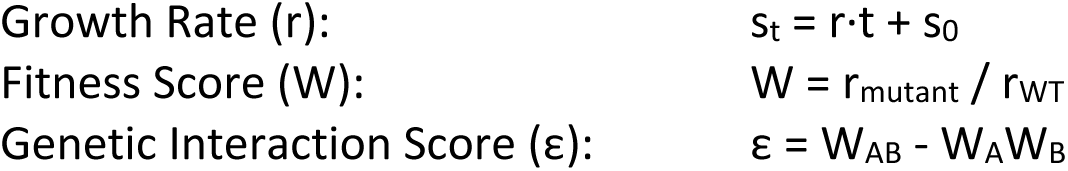

W_AB_ = fitness score for the double-mutant progeny, W_A_ = fitness score for the *MAT***a** parent, W_B_ = fitness score for the *MAT*α mutant, s_t_ = colony size at time t, and s_0_ = colony size at time 0

A histogram binning procedure was used to estimate the mode for genetic interaction scores across biological replicates (up to 20 independent crosses). The “consensus” GI score reported in Figure 5 and Table 3 is the midpoint of the bin containing the maximum number of values (additional details in the supplement).

## Acknowledgements

This work was supported by NIH grant GM078989 and NSF Award #1759900

## Supporting Information Legends

**Figure S1. Example of plate images and growth curves. (A)** Unprocessed (left) and processed (right) images of a phenotyping plate across the 5 time points. **(B)** Growth curves for one of the quadruplicates shown in A before (top) and after (bottom) normalization. Each colored line represents one of the colonies in the quadruplicate. The black and blue lines plot edge colonies (positions C1 and D1), while the red and green lines plot non-edge colonies (C2 and D2).

**Figure S2. Normalizations for six representative phenotyping plates.** Normalizations based on the mean growth rate of the wildtype controls on each plate were used to account for edge effects. Heat maps show a visual representation of growth rates across each plate. The X and Y axis are the coordinates for the 384 positions where a colony may appear. In every case, wild-type controls are in rows A, B, I, and J, columns 1-4, 11-14, and 21-24. Histograms compare the growth rate of colonies that are on the edge of the plate or adjacent to an empty position (distance level 0) with those that are one or more positions away from an edge (non-zero distance levels). The p-value reported above the histogram marks the significance of the difference between the growth rate of edge-adjacent colonies and internal colonies. In each case, raw, unnormalized heat maps, histograms, and p-values are shown just above their normalized counterparts.

**Figure S3. Comparison of fitness scores for double mutants in all four sets of crosses on YPR media.** White cells indicate zero growth and grey cells indicate missing or excluded data. Royal blue is used to designate fitness scores that differ from WT by fewer than 2 standard deviations. Cyan and green indicate fitness scores that are greater than WT by up to or more than 6 standard deviations respectively. Magenta and red indicate fitness scores that are less than WT by up to or more than 6 standard deviations respectively. **A & B)** Cross 1. **C & D)** Cross 2. **E & F)** Cross 3. **G & H)** Cross 4.

**Figure S4. Comparison of fitness scores for double mutants in all four sets of crosses on YPG media.** White cells indicate zero growth and grey cells indicate missing or excluded data. Royal blue is used to designate fitness scores that differ from WT by fewer than 2 standard deviations. Cyan and green indicate fitness scores that are greater than WT by up to or more than 6 standard deviations respectively. Magenta and red indicate fitness scores that are less than WT by up to or more than 6 standard deviations respectively. **A & B)** Cross 1. **C & D)** Cross 2. **E & F)** Cross 3. **G & H)** Cross 4.

**Figure S5. Comparison of fitness scores for double mutants in all four sets of crosses on YPD-Ben media.** White cells indicate zero growth and grey cells indicate missing or excluded data. Royal blue is used to designate fitness scores that differ from WT by fewer than 2 standard deviations. Cyan and green indicate fitness scores that are greater than WT by up to or more than 6 standard deviations respectively. Magenta and red indicate fitness scores that are less than WT by up to or more than 6 standard deviations respectively. **A & B)** Cross 1. **C & D)** Cross 2. **E & F)** Cross 3. **G & H)** Cross 4.

**Table S1.** Parent Strains used in this study

**Table S2.** Primers used in this study

**Table S3.** Potential sources of false negatives

**Table S4.** Replicates supporting synthetic lethal (SL)

**Table S5.** Colony size and growth rate variability across biological replicates of *cdh1Δ swi4Δ*

**Table S6.** Mann-Whitney tests before and after edge effect normalization

**Table S7.** Basal parameter values for wild-type cells. Adjusted parameters are highlighted

**Additional Data Figure 1:** Halo assays of single mutant parents and double mutant progeny for the *MAT***a** kan x *MAT*α SGA nat cross (parent set 1 x parent set 8)

**Additional Data Figure 2:** Halo assays of single mutant parents and double mutant progeny for the *MAT***a** nat x *MAT*α SGA kan cross (parent set 2 x parent set 7)

**Additional Data Figure 3**: Halo assays of single mutant parents and double mutant progeny for the *MAT***a** SGA kan x *MAT*α nat cross (parent set 5 x parent set 4)

**Additional Data Figure 4:** Halo assays of single mutant parents and double mutant progeny for the *MAT***a** SGA nat x *MAT*α kan cross (parent set 3 x parent set 6)

**Additional Data Figure 5:** Comparison of double mutant heterozygous diploids with *MAT***a** and *MAT*α haploid progeny for the *MAT***a** kan x *MAT*α SGA nat cross (parent set 1 x parent set 8)

**Additional Data Figure 6:** Comparison of double mutant heterozygous diploids with *MAT***a** and *MAT*α haploid progeny for the *MAT***a** nat x *MAT*α SGA kan cross (parent set 2 x parent set 7)

**Additional Data Figure 7:** Comparison of double mutant heterozygous diploids with *MAT***a** and MATalpha haploid progeny for the *MAT***a** SGA kan x *MAT*α nat cross (parent set 5 x parent set 4)

**Additional Data Figure 8:** Comparison of double mutant heterozygous diploids with *MAT***a** and *MAT*α haploid progeny for the *MAT***a** SGA nat x *MAT*α kan cross (parent set 3 x parent set 6)

**Additional Data Table 1:** Colony size data for all double mutant haploid progeny selected for on YPD+G418/nat.

**Additional Data Table 2:** Growth rates for single mutant parents on all media types

**Additional Data Table 3:** Growth rates for *MAT***a** and *MAT*α double mutant progeny from the *MAT***a** kan x *MAT*α SGA nat cross (parent set 1 x parent set 8) on all media types

**Additional Data Table 4:** Growth rates for *MAT***a** and *MAT*α double mutant progeny from the *MAT***a** nat x *MAT*α SGA kan cross (parent set 2 x parent set 7) on all media types

**Additional Data Table 5:** Growth rates for *MAT***a** and *MAT*α double mutant progeny from the *MAT***a** SGA kan x *MAT*α nat cross (parent set 5 x parent set 4) on all media types

**Additional Data Table 6:** Growth rates for *MAT***a** and *MAT*α double mutant progeny from the *MAT***a** SGA nat x *MAT*α kan cross (parent set 3 x parent set 6) on all media types

**Additional Data Table 7:** Fitness scores for single mutant parents on all media types

**Additional Data Table 8:** Fitness scores for *MAT***a** and *MAT*α double mutant progeny from the *MAT***a** kan x *MAT*α SGA nat cross (parent set 1 x parent set 8) on all media types

**Additional Data Table 9:** Fitness scores for *MAT***a** and *MAT*α double mutant progeny from the *MAT***a** nat x *MAT*α SGA kan cross (parent set 2 x parent set 7) on all media types

**Additional Data Table 10:** Fitness scores for *MAT***a** and *MAT*α double mutant progeny from the *MAT***a** SGA kan x *MAT*α nat cross (parent set 5 x parent set 4) on all media types

**Additional Data Table 11:** Fitness scores for *MAT***a** and *MAT*α double mutant progeny from the *MAT***a** SGA nat x *MAT*α kan cross (parent set 3 x parent set 6) on all media types

**Additional Data Table 12:** Genetic interaction scores for double mutant progeny from all crosses on all media types. Monogenic crosses and crosses where a parent was not generated or had a fitness score of zero are marked NA.

## References

1. Morgan, D. & Morgan, D.O. The Cell Cycle: Principles of Control. (OUP/New Science Press, 2007).

2. Csikasz-Nagy, A., Palmisano, A. & Zamborszky, J. Molecular network dynamics of cell cycle control: transitions to start and finish. Methods Mol Biol 761, 277–291 (2011).

3. Alberghina, L., Hofer, T. & Vanoni, M. Molecular networks and system-level properties. J Biotechnol 144, 224–233 (2009).

4. Wen, Z. et al. Identifying responsive modules by mathematical programming: an application to budding yeast cell cycle. PLoS One 7, e41854 (2012).

5. Ideker, T.E. Network genomics. Ernst Schering Res Found Workshop, 89–115 (2007).

6. Carter, G.W. et al. A systems-biology approach to modular genetic complexity. Chaos 20, 026102 (2010).

7. Ramon, C., Gollub, M.G. & Stelling, J. Integrating-omics data into genome-scale metabolic network models: principles and challenges. Essays Biochem 62, 563–574 (2018).

8. Trescher, S., Munchmeyer, J. & Leser, U. Estimating genome-wide regulatory activity from multi-omics data sets using mathematical optimization. BMC Syst Biol 11, 41 (2017).

9. Buescher, J.M. & Driggers, E.M. Integration of omics: more than the sum of its parts. Cancer Metab 4, 4 (2016).

10. Bersanelli, M. et al. Methods for the integration of multi-omics data: mathematical aspects. BMC Bioinformatics 17 **Suppl 2**, 15 (2016).

11. Yilmaz, L.S. & Walhout, A.J. Metabolic network modeling with model organisms. Curr Opin Chem Biol 36, 32–39 (2017).

12. Hou, J., Acharya, L., Zhu, D. & Cheng, J. An overview of bioinformatics methods for modeling biological pathways in yeast. Brief Funct Genomics 15, 95–108 (2016).

13. Sanchez, B.J. & Nielsen, J. Genome scale models of yeast: towards standardized evaluation and consistent omic integration. Integr Biol (Camb) 7, 846–858 (2015).

14. Garcia-Campos, M.A., Espinal-Enriquez, J. & Hernandez-Lemus, E. Pathway Analysis: State of the Art. Front Physiol 6, 383 (2015).

15. Brodland, G.W. How computational models can help unlock biological systems. Semin Cell Dev Biol 47-48, 62–73 (2015).

16. Soh, K.C., Miskovic, L. & Hatzimanikatis, V. From network models to network responses: integration of thermodynamic and kinetic properties of yeast genome-scale metabolic networks. FEMS Yeast Res 12, 129–143 (2012).

17. Alberghina, L., Coccetti, P. & Orlandi, I. Systems biology of the cell cycle of Saccharomyces cerevisiae: From network mining to system-level properties. Biotechnol Adv 27, 960–978 (2009).

18. Barberis, M., Todd, R.G. & van der Zee, L. Advances and challenges in logical modeling of cell cycle regulation: perspective for multi-scale, integrative yeast cell models. FEMS Yeast Res 17 (2017).

19. Kraikivski, P., Chen, K.C., Laomettachit, T., Murali, T.M. & Tyson, J.J. From START to FINISH: computational analysis of cell cycle control in budding yeast. NPJ Syst Biol Appl 1, 15016 (2015).

20. Radmaneshfar, E. et al. From START to FINISH: the influence of osmotic stress on the cell cycle. PLoS One 8, e68067 (2013).

21. Barik, D., Ball, D.A., Peccoud, J. & Tyson, J.J. A Stochastic Model of the Yeast Cell Cycle Reveals Roles for Feedback Regulation in Limiting Cellular Variability. PLOS Computational Biology 12, e1005230 (2016).

22. Chen, K.C. et al. Integrative analysis of cell cycle control in budding yeast. Mol Biol Cell 15, 3841–3862 (2004).

23. Ball, D.A. et al. Stochastic exit from mitosis in budding yeast: model predictions and experimental observations. Cell cycle (Georgetown, Tex 10, 999–1009 (2011).

24. Ball, D.A. et al. Oscillatory Dynamics of Cell Cycle Proteins in Single Yeast Cells Analyzed by Imaging Cytometry. PLoS ONE 6, 12 (2011).

25. Ball, D.A. et al. Measurement and modeling of transcriptional noise in the cell cycle regulatory network. Cell cycle (Georgetown, Tex 12, 3203–3218 (2013).

26. Adames, N.R. et al. Experimental testing of a new integrated model of the budding yeast START transition. Molecular Biology of the Cell (2015).

27. Alberghina, L., Rossi, R.L., Wanke, V., Querin, L. & Vanoni, M. Checking cell size in budding yeast: a systems biology approach. Ital J Biochem 52, 55–57 (2003).

28. Adames, N.R., Gallegos, J.E. & Peccoud, J. Yeast genetic interaction screens in the age of CRISPR/Cas. Curr Genet 65, 307–327 (2019).

29. Tong, A.H. & Boone, C. Synthetic genetic array analysis in Saccharomyces cerevisiae. Methods Mol Biol 313, 171–192 (2006).

30. Schuldiner, M. et al. Exploration of the function and organization of the yeast early secretory pathway through an epistatic miniarray profile. Cell 123, 507–519 (2005).

31. Ikui, A.E. & Cross, F.R. Specific genetic interactions between spindle assembly checkpoint proteins and B-Type cyclins in Saccharomyces cerevisiae. Genetics 183, 51–61 (2009).

32. Proudfoot, K.G. et al. Checkpoint Proteins Bub1 and Bub3 Delay Anaphase Onset in Response to Low Tension Independent of Microtubule-Kinetochore Detachment. Cell Reports 27, 416–428.e414 (2019).

33. Adames, N.R. et al. Experimental testing of a new integrated model of the budding yeast Start transition. Mol Biol Cell 26, 3966–3984 (2015).

34. Daniel, J.A., Yoo, J., Bettinger, B.T., Amberg, D.C. & Burke, D.J. Eliminating gene conversion improves high-throughput genetics in Saccharomyces cerevisiae. Genetics 172, 709–711 (2006).

35. Gallegos, J., N. Adames, S. Hayrynen, J. Peccoud Challenges and opportunities for strain verification by whole-genome sequencing. bioRxiv, 515338 (2019).

36. Singh, I., Pass, R., Togay, S.O., Rodgers, J.W. & Hartman, J.L.t. Stringent mating-type-regulated auxotrophy increases the accuracy of systematic genetic interaction screens with Saccharomyces cerevisiae mutant arrays. Genetics 181, 289–300 (2009).

37. Lesage, G. et al. Analysis of beta-1,3-glucan assembly in Saccharomyces cerevisiae using a synthetic interaction network and altered sensitivity to caspofungin. Genetics 167, 35–49 (2004).

38. Tong, A.H. et al. Global mapping of the yeast genetic interaction network. Science 303, 808–813 (2004).

39. Vizeacoumar, F.J. et al. Integrating high-throughput genetic interaction mapping and high-content screening to explore yeast spindle morphogenesis. The Journal of cell biology 188, 69–81 (2010).

40. Tong, A.H. et al. Systematic genetic analysis with ordered arrays of yeast deletion mutants. Science 294, 2364–2368 (2001).

41. Sarin, S. et al. Uncovering novel cell cycle players through the inactivation of securin in budding yeast. Genetics 168, 1763–1771 (2004).

42. St Onge, R.P., et al. Systematic pathway analysis using high-resolution fitness profiling of combinatorial gene deletions. Nat Genet 39, 199–206 (2007).

43. Bandyopadhyay, S. et al. Rewiring of genetic networks in response to DNA damage. Science 330, 1385–1389 (2010).

44. Baryshnikova, A. et al. Global linkage map connects meiotic centromere function to chromosome size in budding yeast. G3 (Bethesda) 3, 1741–1751 (2013).

45. Wagih, O. et al. SGAtools: one-stop analysis and visualization of array-based genetic interaction screens. Nucleic Acids Res 41, W591–596 (2013).

46. Kuzmin, E., Costanzo, M., Andrews, B. & Boone, C. Synthetic Genetic Array Analysis. Cold Spring Harb Protoc 2016, pdb prot088807 (2016).

47. Wheals, B.B. Simple preparation of a bonded cation-exchange packing material and its application to the separation of phenothiazines by high-performance liquid chromatography. J Chromatogr 177, 263–270 (1979).

48. Barik, D., Ball, D.A., Peccoud, J. & Tyson, J.J. A Stochastic Model of the Yeast Cell Cycle Reveals Roles for Feedback Regulation in Limiting Cellular Variability. PLoS Comput Biol 12, e1005230 (2016).

49. Li, R. & Murray, A.W. Feedback control of mitosis in budding yeast. Cell 66, 519–531 (1991).

50. Dudley, A.M., Janse, D.M., Tanay, A., Shamir, R. & Church, G.M. A global view of pleiotropy and phenotypically derived gene function in yeast. Mol Syst Biol 1, 2005 0001 (2005).

51. Parsons, A.B. et al. Integration of chemical-genetic and genetic interaction data links bioactive compounds to cellular target pathways. Nat Biotechnol 22, 62–69 (2004).

52. Brown, J.A. et al. Global analysis of gene function in yeast by quantitative phenotypic profiling. Mol Syst Biol 2, 2006 0001 (2006).

53. Chen, S.H., Albuquerque, C.P., Liang, J., Suhandynata, R.T. & Zhou, H. A proteome-wide analysis of kinase-substrate network in the DNA damage response. J Biol Chem 285, 12803–12812 (2010).

54. Duffy, S. et al. Overexpression screens identify conserved dosage chromosome instability genes in yeast and human cancer. Proc Natl Acad Sci U S A 113, 9967–9976 (2016).

55. Kapitzky, L. et al. Cross-species chemogenomic profiling reveals evolutionarily conserved drug mode of action. Mol Syst Biol 6, 451 (2010).

56. Shively, C.A. et al. Genetic networks inducing invasive growth in Saccharomyces cerevisiae identified through systematic genome-wide overexpression. Genetics 193, 1297–1310 (2013).

57. Wang, S.H. et al. Curcumin-Mediated HDAC Inhibition Suppresses the DNA Damage Response and Contributes to Increased DNA Damage Sensitivity. PLoS One 10, e0134110 (2015).

58. Woolstencroft, R.N. et al. Ccr4 contributes to tolerance of replication stress through control of CRT1 mRNA poly(A) tail length. J Cell Sci 119, 5178–5192 (2006).

59. Wu, X. & Jiang, Y.W. Genetic/genomic evidence for a key role of polarized endocytosis in filamentous differentiation of S. cerevisiae. Yeast 22, 1143–1153 (2005).

60. Iouk, T., Kerscher, O., Scott, R.J., Basrai, M.A. & Wozniak, R.W. The yeast nuclear pore complex functionally interacts with components of the spindle assembly checkpoint. J Cell Biol 159, 807–819 (2002).

61. Tatchell, K., Makrantoni, V., Stark, M.J. & Robinson, L.C. Temperature-sensitive ipl1-2/Aurora B mutation is suppressed by mutations in TOR complex 1 via the Glc7/PP1 phosphatase. Proc Natl Acad Sci U S A 108, 3994–3999 (2011).

62. Daniel, J.A., Keyes, B.E., Ng, Y.P., Freeman, C.O. & Burke, D.J. Diverse functions of spindle assembly checkpoint genes in Saccharomyces cerevisiae. Genetics 172, 53–65 (2006).

63. Doncic, A., Ben-Jacob, E., Einav, S. & Barkai, N. Reverse engineering of the spindle assembly checkpoint. PLoS One 4, e6495 (2009).

64. Choi, J.E. & Chung, W.H. Synthetic lethal interaction between oxidative stress response and DNA damage repair in the budding yeast and its application to targeted anticancer therapy. J Microbiol 57, 9–17 (2019).

65. Diaz-Mejia, J.J. et al. Mapping DNA damage-dependent genetic interactions in yeast via party mating and barcode fusion genetics. Mol Syst Biol 14, e7985 (2018).

66. Leung, G.P., Aristizabal, M.J., Krogan, N.J. & Kobor, M.S. Conditional genetic interactions of RTT107, SLX4, and HRQ1 reveal dynamic networks upon DNA damage in S. cerevisiae. G3 (Bethesda) 4, 1059–1069 (2014).

67. Donovan, J.D., Toyn, J.H., Johnson, A.L. & Johnston, L.H. P40SDB25, a putative CDK inhibitor, has a role in the M/G1 transition in Saccharomyces cerevisiae. Genes Dev 8, 1640–1653 (1994).

68. Schwob, E., Böhm, T., Mendenhall, M.D. & Nasmyth, K. The B-type cyclin kinase inhibitor p40SIC1 controls the G1 to S transition in S. cerevisiae. Cell 79, 233–244 (1994).

69. Nugroho, T.T. & Mendenhall, M.D. An inhibitor of yeast cyclin-dependent protein kinase plays an important role in ensuring the genomic integrity of daughter cells. Molecular and Cellular Biology 14, 3320 (1994).

70. Sprague, G.F., Jr. Assay of yeast mating reaction. Methods Enzymol 194, 77–93 (1991).

71. Hoyt, M.A., Totis, L. & Roberts, B.T. S. cerevisiae genes required for cell cycle arrest in response to loss of microtubule function. Cell 66, 507–517 (1991).

72. Silva, P. et al. Monitoring the fidelity of mitotic chromosome segregation by the spindle assembly checkpoint. Cell Prolif 44, 391–400 (2011).

73. Eng, W.K., Faucette, L., Johnson, R.K. & Sternglanz, R. Evidence that DNA topoisomerase I is necessary for the cytotoxic effects of camptothecin. Mol Pharmacol 34, 755–760 (1988).

74. Kjeldsen, E., Svejstrup, J.Q., Gromova, II, Alsner, J. & Westergaard, O. Camptothecin inhibits both the cleavage and religation reactions of eukaryotic DNA topoisomerase I. J Mol Biol 228, 1025–1030 (1992).

75. Turner, M.K., Abrams, R. & Lieberman, I. Meso-alpha, beta-diphenylsuccinate and hydroxyurea as inhibitors of deoxycytidylate synthesis in extracts of Ehrlich ascites and L cells. J Biol Chem 241, 5777–5780 (1966).

76. Cortez, D. Preventing replication fork collapse to maintain genome integrity. DNA Repair (Amst) 32, 149–157 (2015).

77. Martin-Yken, H. et al. Saccharomyces cerevisiae YCRO17c/CWH43 encodes a putative sensor/transporter protein upstream of the BCK2 branch of the PKC1-dependent cell wall integrity pathway. Yeast 18, 827–840 (2001).

78. Wijnen, H. & Futcher, B. Genetic analysis of the shared role of CLN3 and BCK2 at the G(1)-S transition in Saccharomyces cerevisiae. Genetics 153, 1131–1143 (1999).

79. Epstein, C.B. & Cross, F.R. Genes that can bypass the CLN requirement for Saccharomyces cerevisiae cell cycle START. Mol Cell Biol 14, 2041–2047 (1994).

80. Di Como, C.J., Chang, H. & Arndt, K.T. Activation of CLN1 and CLN2 G1 cyclin gene expression by BCK2. Mol Cell Biol 15, 1835–1846 (1995).

81. Bastajian, N., Friesen, H. & Andrews, B.J. Bck2 acts through the MADS box protein Mcm1 to activate cell-cycle-regulated genes in budding yeast. PLoS Genet 9, e1003507 (2013).

82. Jorgensen, P., Nishikawa, J.L., Breitkreutz, B.J. & Tyers, M. Systematic identification of pathways that couple cell growth and division in yeast. Science 297, 395–400 (2002).

83. Geymonat, M., Spanos, A., de Bettignies, G. & Sedgwick, S.G. Lte1 contributes to Bfa1 localization rather than stimulating nucleotide exchange by Tem1. J Cell Biol 187, 497–511 (2009).

84. Pan, X. et al. A DNA integrity network in the yeast Saccharomyces cerevisiae. Cell 124, 1069–1081 (2006).

85. Reiter, W. et al. Yeast protein phosphatase 2A-Cdc55 regulates the transcriptional response to hyperosmolarity stress by regulating Msn2 and Msn4 chromatin recruitment. Mol Cell Biol 33, 1057–1072 (2013).

86. Schwab, M., Lutum, A.S. & Seufert, W. Yeast Hct1 is a regulator of Clb2 cyclin proteolysis. Cell 90, 683–693 (1997).

87. Archambault, V. et al. Genetic and biochemical evaluation of the importance of Cdc6 in regulating mitotic exit. Mol Biol Cell 14, 4592–4604 (2003).

88. Cross, F.R., Yuste-Rojas, M., Gray, S. & Jacobson, M.D. Specialization and targeting of B-type cyclins. Mol Cell 4, 11–19 (1999).

89. Surana, U. et al. The role of CDC28 and cyclins during mitosis in the budding yeast S. cerevisiae. Cell 65, 145–161 (1991).

90. Richardson, H., Lew, D.J., Henze, M., Sugimoto, K. & Reed, S.I. Cyclin-B homologs in Saccharomyces cerevisiae function in S phase and in G2. Genes Dev 6, 2021–2034 (1992).

91. Pecani, K. & Cross, F.R. Degradation of the Mitotic Cyclin Clb3 Is not Required for Mitotic Exit but Is Necessary for G_1_ Cyclin Control of the Succeeding Cell Cycle. Genetics 204, 1479–1494 (2016).

92. Wasch, R. & Cross, F.R. APC-dependent proteolysis of the mitotic cyclin Clb2 is essential for mitotic exit. Nature 418, 556–562 (2002).

93. Cross, F.R., Schroeder, L. & Bean, J.M. Phosphorylation of the Sic1 inhibitor of B-type cyclins in Saccharomyces cerevisiae is not essential but contributes to cell cycle robustness. Genetics 176, 1541–1555 (2007).

94. Queralt, E. & Igual, J.C. Functional connection between the Clb5 cyclin, the protein kinase C pathway and the Swi4 transcription factor in Saccharomyces cerevisiae. Genetics 171, 1485–1498 (2005).

95. Ye, P. et al. Gene function prediction from congruent synthetic lethal interactions in yeast. Mol Syst Biol 1, 2005 0026 (2005).

96. Koch, C., Moll, T., Neuberg, M., Ahorn, H. & Nasmyth, K. A role for the transcription factors Mbp1 and Swi4 in progression from G1 to S phase. Science 261, 1551–1557 (1993).

97. Queralt, E. & Igual, J.C. Cell cycle activation of the Swi6p transcription factor is linked to nucleocytoplasmic shuttling. Mol Cell Biol 23, 3126–3140 (2003).

98. Alepuz, P.M., Matheos, D., Cunningham, K.W. & Estruch, F. The Saccharomyces cerevisiae RanGTP-binding protein msn5p is involved in different signal transduction pathways. Genetics 153, 1219–1231 (1999).

99. Toone, W.M. et al. Rme1, a negative regulator of meiosis, is also a positive activator of G1 cyclin gene expression. EMBO J 14, 5824–5832 (1995).

100. Breeden, L. & Mikesell, G. Three independent forms of regulation affect expression of HO, CLN1 and CLN2 during the cell cycle of Saccharomyces cerevisiae. Genetics 138, 1015–1024 (1994).

