## Supplement for "Genetic interactions derived from high-throughput phenotyping of 7,350 yeast cell cycle mutants"

### Parent Strain Construction

We used several strategies to generate the eight sets of parent strains used in this study (Table S1). We obtained most of the Set 1 and Set 3 strains by sporulation and tetrad analysis of the heterozygous diploid commercial collection of *kanMX* strains [1, 2], but we made some by *de novo kanMX* PCR-mediated gene deletions in BY4741 or BY4742 (using pFA6a-kanMX as the template and listed primers[3-6], Table S2). We obtained most of the Set 2 and Set 4 strains by transformation of the heterozygous diploid strains with a *natMX* PCR product (using MX.for and MX.rev primers in Table S2 with pAG25 template. pAG25 and its sequences are available from Addgene) to switch markers, followed by selection of nourseothricin-resistant/G418-sensitive transformants, sporulation, and tetrad analysis. We made the rest of sets 2 and 4 by *de novo natMX* PCR-mediated gene deletions in BY4741 or BY4742 (using pAG25 as the template and listed primers). We made most of Set 7 and Set 8 by *de novo* PCR-mediated gene deletions in the SGA strain Y8205 [7], but we made some of these strains by crossing one of the BY4741-derived gene deletion strain with Y8205, followed by tetrad dissection. We then used the SGA method to cross these strains to BY4741 and obtain *MAT***a** versions of these strains for Set 5 and Set 6. All strains were confirmed by PCR of genomic DNA using one set of test primers for the gene deletion and another for the wild-type gene[8] (Table S2). All strains (parents and progeny) are available upon request.

### Double Mutant Progeny Construction

All crosses followed a standard format in which the *MAT*$\alpha$ strains (Sets 3, 4, 7, and 8) which we will call the "hit" strains, were arrayed alphabetically by gene name so that each strain was a single well in a 36-well block, with two replicate *MAT*$\alpha$ blocks per plate, leaving the first and fifth rows empty for the addition of the wild-type parents during phenotyping. If a deletion strain was missing in a *MAT*$\alpha$ set, we left the position empty. We arrayed the *MAT***a** strains, which we will call the query or "bait" strains, so that each *MAT***a** strain in the set fills a block of 36 wells at the same positions as one of the two blocks of *MAT*$\alpha$ strains (i.e., in rows 2-4 or 6-8).

Before crossing, each set of parent strains was arrayed in 96-well microtiter PlusPlates (Singer Instruments, Somerset, UK) containing YPD broth and pinned onto YPD+G418 (300 µg/ml; odd numbered sets) or YPD+nat (150 µg/ml; even numbered sets) and grown for 3-4 days at 30 °C.

For the crosses, we used a Rotor HDA (Singer Instruments, Somerset, UK) to replica-pin each *MAT*$\alpha$ plate to 12-18 YPD plates using 96 long repads with 6 wet mix cycles and 4 dry mix cycles to ensure robust inoculation of each plate. Visual inspection of each plate ensured proper transfer of cells. We then pinned each *MAT***a** plate on top of one *MAT*$\alpha$ plate using the same conditions to ensure good mixing of the two parent strains on the YPD plate. Matings were performed on YPD at 30 °C for two days.

Diploids were selected on YPD + G418/nat (300/150 µg/ml) at 30 °C for two days. Diploids were sporulated on enriched sporulation media (1% potassium acetate, 0.1% yeast extract, 0.05% glucose, 0.1 g of his/leu/lys/ura supplement) at 24 °C for five days.

Haploid progeny were selected as described in Tong et al. (2006) [7], except that we separately selected for both *MAT***a** and *MAT*$\alpha$ progeny. We first selected haploids from the sporulation plates by replica-pinning on SD-arg/his/lys+canavanine/thialysine (100/100 µg/ml) for *MAT***a** progeny and SD-arg/leu/lys+canavanine/thialysine (100 µg/ml) for *MAT*$\alpha$ progeny. For the second round of haploid selection, we added G418 (300 µg/ml) to these plates. For the final haploid selection, we added both G418 (600 µg/ml) and nourseothricin (150 µg/ml) to obtain double mutant haploids.

### Halo Assays

Halo assays were used to confirm the mating type of the parents and progeny and identify potential cell signaling and chromosome segregation defects**.** We performed halo assays as described[9] using Y955 lawns to test for **a**-halos, and Y991 to test for $\alpha$-halos. We prepared halo assay plates by growing Y995 and Y991 in YPD broth at 30 °C overnight in a shaking incubator. The next morning, we diluted each strain 1/5 in YPD broth, vortexed the tubes, and used sterile glass beads to spread 500 µl of the dilution per plate onto YPD PlusPlates. We allowed these plates to dry before pinning each plate of parent strains or haploid progeny (at 96 colony densities) onto both a Y995 lawn (to test for **a**-factor secretion) and a Y991 lawn (to test for $\alpha$-factor secretion). We imaged halo assay plates after 48 h of growth (see Additional Data). Individual lines that did not behave as expected (most likely due to isolated genetic mishaps) were not excluded, as the consensus between multiple biological replicates was used to draw the conclusions presented.

### Identifying Curation Errors

For several of the manually curated synthetic lethal interactions on the Saccharomyces Genome Database, we found that the listed SL interaction was not supported by the cited paper. In some cases, what was curated as synthetic lethal was in fact not lethal but exhibited some other kind of growth defect. In other cases, the interaction was lethal, but was tested in a mutant background in which one or more additional cell cycle genes were knocked out. Those synthetic lethal interactions that we found to be curation errors are listed, along with their references, below:

*clb5Δ clb6Δ*[10-12], *cdc55Δ cln1Δ*[13], *cdc55Δ cln2Δ*[13], *cln1Δ cln2Δ*[13-19], *cln1Δ cln3Δ*[16, 17], *cln1Δ msn5Δ*[20], *cln2Δ cln3Δ*[16, 17], *cln2Δ msn5Δ*[20], *fkh1Δ fkh2Δ*[21], *ssa1Δ ydj1Δ*[22]

### Tetrad Analysis

To identify synthetic lethality by tetrad analysis, several tetrads (usually 12) were dissected for one or more biological replicates of each of the 58 gene combinations listed in Table 2. The surviving spores were patched onto YPD plates, and then replica plated onto YPD+G418 (600ug/ml), YPD+nat (150ug/ml), and Y995 and Y991 lawn plates (as described above but using 300ul diluted culture).

Dissections from which we could recover very few live spores of any genotype were identified as having a likely meiotic defect (MD). Recovery of a live spore that was resistant to both antibiotics was considered evidence for viability (V). Cases where a dead spore in a tetrad could be inferred to have the double mutant phenotype based on allele segregation were considered evidence for synthetic lethality (SL). If the ratio of spores supporting synthetic lethality to spores supporting viability (SL:V) was 4:1 or greater, we considered the gene combination SL. If the SL:V ratio was between 4:1 and 1:1, we considered the gene combination to have reduced viability (RV). If the SL:V ratio was less than 1:1, we considered the gene combination to be viable (V). Cases where we were able to identify fewer than two spores as potentially SL or V (due to low viability overall) were also designated MD.

### Data Analysis

#### Growth rates

To account for plate-to-plate variation and edge effects, we normalized all colonies on a plate-by-plate basis using in-plate controls consisting of six sets of BY4741 and BY4742 wild-type parent strains in various positions on the plate (Figure 1). Within plate spatial effects are probably negligible in our case, because the Rotor HDA system uses disposable pin pads with fixed pins (rather than floating pins). Consequently, differences in plate thickness result in obvious failures to evenly transfer colonies (or complete failure to transfer) in certain areas of the plate. After visual inspection of all plates, we disposed of failed plates and repeated the replica-pinning to obtain consistent colony transfer across the plate.

We estimated growth rates for all wild-type, parent and progeny strains using the same procedure (Figure S1). For each position on a plate, we used a simple linear model to estimate the growth rate based on the colony sizes measured from 0 to 60 hours. Any positions having a zero colony size at any time point were discarded. For the remaining positions we estimated growth rate using a simple linear model to predict colony size as a function of time:

$$s_t = r \cdot t + s_0$$

Here, $s_t$ is the colony size at time $t$, $r$ is the growth rate in pixels/hour and $s_0$ is the colony size at time 0. Note that the bias term $s_0$ accounts for variation in the initial colony sizes. In rare cases a colony would exhibit a negative growth rate, in which case we discarded that position, assuming experimental error. Examples of colony sizes and growth rates for the combination *cdh1Δ swi5Δ* are provided in Table S5.

We tested growth rates for "edge effects" that could arise either from colonies growing adjacent to plate edges, or those adjacent to empty positions (dark areas in Figure S2). Because of these additional empty positions, we categorized all positions on a plate based on their distance from an empty (zero growth) colony. Our hypothesis was that a zero-growth colony, like an edge, may supply extra growth medium to neighboring positions and hence increase their growth rates.

To account for this, we arbitrarily assign a distance of 0 for positions along an edge or having at least one zero-growth neighbor. All other positions are assigned a distance of 1 greater than their lowest neighbor. We note that this approach does not distinguish between edge positions having several zero-growth neighbors and those having just one. We selected this model for its simplicity; because in practice it adequately addresses variability between distance categories (for examples, see Figure S1 and Figure S2), and because variability among wild-type edge positions tended to be relatively low regardless of the number of zero-growth neighbors.

To test the hypothesis that edge effects might influence our results, we compared the distributions of growth rates for each distance category. We used a Mann-Whitney test to compare these distributions and found that in most cases the differences were significant, even among wild-type colonies (see Table S6). To account for these edge effects, we computed the average growth rate for each distance category, $r_d$, and the average for a whole plate, $r_p$ (always omitting any zero-growth colonies). We then adjusted individual colony growth rates $r_c$ as follows:

$$r_c = r_c \cdot r_p / r_d$$

These adjustments tended to yield distributions with no significant differences at distinct edge distances.

All strains were grown using four technical replicates (quadruplicates) in adjacent positions on a plate. We applied jack-knife filtering to eliminate potential outliers, but in practice found that only a handful of examples yielded outliers.

#### Fitness scores

We computed the fitness score for each parent mutant and for each double-mutant by dividing the average normalized growth rate across each quadruplicate by the average normalized growth rate for all wild-type colonies on the same plate. Overall values ranged from 0.00 to 2.32 with an average of 0.83.

For the heatmaps shown in Figure 2, we present a visual representation of the fitness scores for all mutant combinations for the YPD growth medium. To find potentially significant differences between fitness scores for mutant combinations and wild-type scores on the same plate, we estimate the mean and standard deviation (SD) for wild-type scores and compute the number of SDs between each mutant score and its associated wild-type score:

$$\Delta_{AB} = (W_{AB} - 1)/s_W,$$

Where $\Delta_{AB}$ is the difference in SDs for mutant combination AB; $W_{AB}$ is the mutant fitness score; $s_W$ is the sample SD of wild-type fitness scores, and 1 is the average wild-type score (by definition). This provides a convenient way to compare the relative significance of fitness scores across different plates. Highlighting noteworthy fitness scores is then a matter of selecting categories based on each score's distance from wild-type, measured in SDs. We establish five different categories: scores within two SDs of wild-type; scores more than two SDs away (higher or lower), and scores more than six SDs away (higher or lower). In practice we had no scores more than six SDs higher than wild-type. We also include categories for combinations exhibiting no growth and for those we discarded.

There were rare cases in which we had duplicate sets of colonies for the same combinations (when the same gene was knocked out with the same marker more than once in a given cross). Erring on the conservative side, we elected to represent the fitness score with the *lowest* SD difference in the figures (Figure 4, and Figure S3-Figure S7).

#### Genetic interaction scores

Having obtained fitness scores for each parent and each double-mutant, we compute the genetic interaction score by:

$$\varepsilon = W_{AB} - W_A W_B$$

Here, $W_{AB}$ is the fitness score for the double-mutant progeny, $W_A$ is the fitness score for the *MAT***a** parent, and $W_B$ is the fitness score for the *MAT*α mutant. Overall genetic interaction scores ranged from -1.4 to 1.2 with an average score of -0.0013.

In the heatmaps shown in Figure 5, and the values shown in Table 3, we summarize the genetic interaction scores for each mutant combination, arriving at a single characteristic score for each. These scores are based on up to 20 distinct measurements depending on the number of colonies retained after filtering. The scores for each combination tend to vary widely across replicates, presumably due to genetic mishaps. Genetic mishaps, such as secondary mutations or nondisjunction will act as outliers, so using mean or median scores can be misleading. Instead we elected to estimate the mode.

The genetic interaction scores are real values, so a list of several values is unlikely to yield a most-common value (mode), since rarely will two values be exactly the same. Instead, we use a histogram binning procedure (the Python numpy library's histogram_bin_edges) to establish *k* equidistant bins based on the distribution of all genetic interaction scores for a given gene combination. The values for each mutant combination are then binned, and we estimate the score as the midpoint of the bin containing the maximum number of values.

In some cases, two or more bins may have the same number of values (see example below). For this reason, we establish sets of bins at various resolutions by using values of *k* from 11 down to 1. For every mutant combination, we attempt to find a maximal bin starting with *k*=11 bins. If two or more bins tie, we next try *k*=10, then *k*=9, and so on. Note that for *k*=1 the resulting score would be the mean of the overall distribution, but in practice the procedure never uses fewer than *k*=4 bins. Cases in which the consensus GI score for each combination were above the top 5% or below the bottom 5% of the distribution of all un-binned scores for a particular growth medium are highlighted in Table 3.

**Example:**

*scores = [-0.31, -0.27, -0.21, -0.17, -0.02, 0.10, 0.20, 0.22, 0.30, 0.30]*

*<u>k=8</u>: [-1.32, -1.02, -0.72, -0.41, -0.11, 0.19, 0.49, 0.79, 1.1]*

*bin counts: [0, 0, 0, 4, 2, 4, 0, 0]; maximal bins are [-0.41,-0.11] and [0.19,0.49]*

*k=7: [-1.32, -0.98, -0.63, -0.29, 0.06, 0.41, 0.75, 1.1]*

*bin counts: [0, 0, 1, 4,* ***5****, 0, 0]; maximal bin is [0.06, 0.41] with midpoint 0.235*

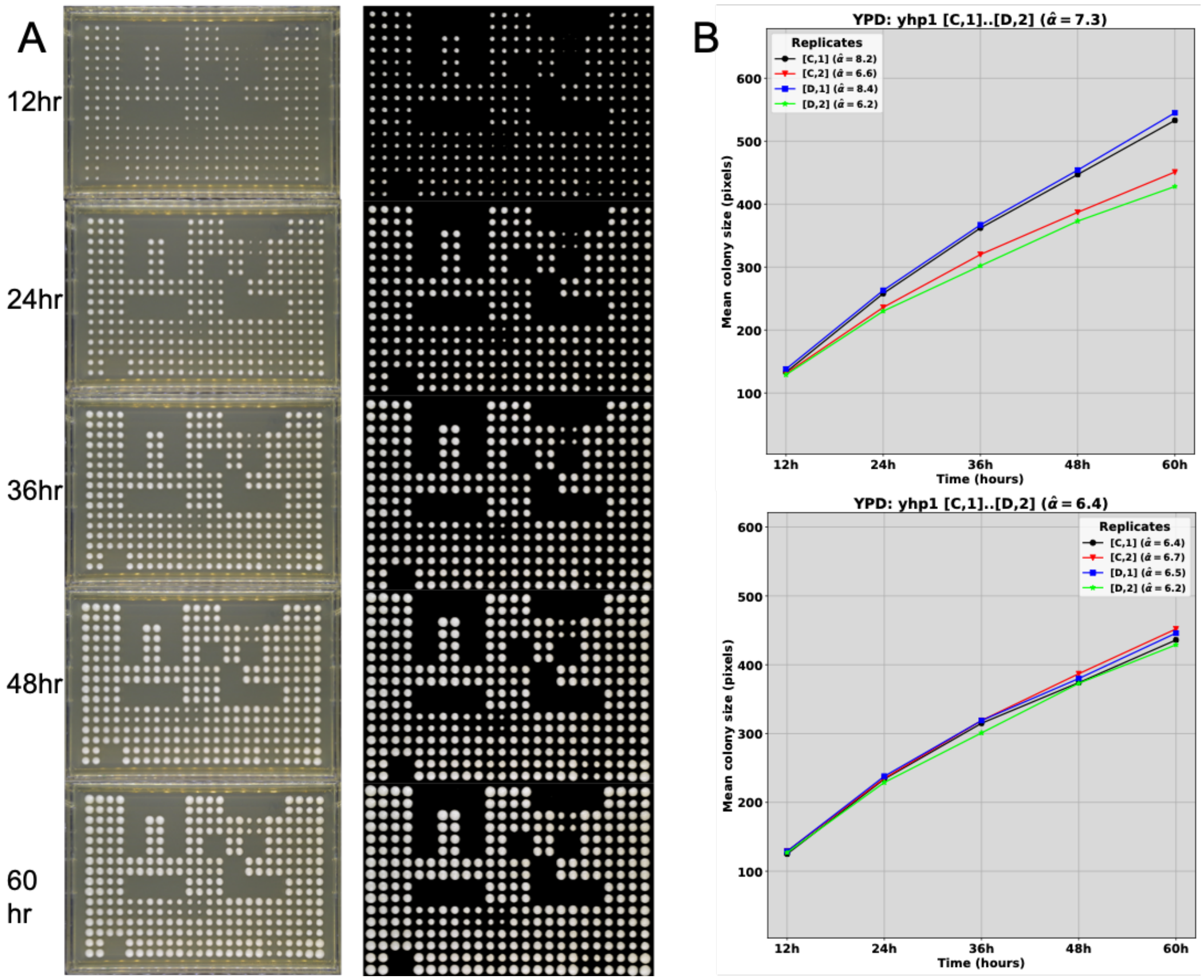


**Figure S1. Example of plate images and growth curves. (A)** Unprocessed (left) and processed (right) images of a phenotyping plate across the 5 time points. **(B)** Growth curves for one of the quadruplicates shown in A before (top) and after (bottom) normalization. Each colored line represents one of the colonies in the quadruplicate. The black and blue lines plot edge colonies (positions C1 and D1), while the red and green lines plot non-edge colonies (C2 and D2).

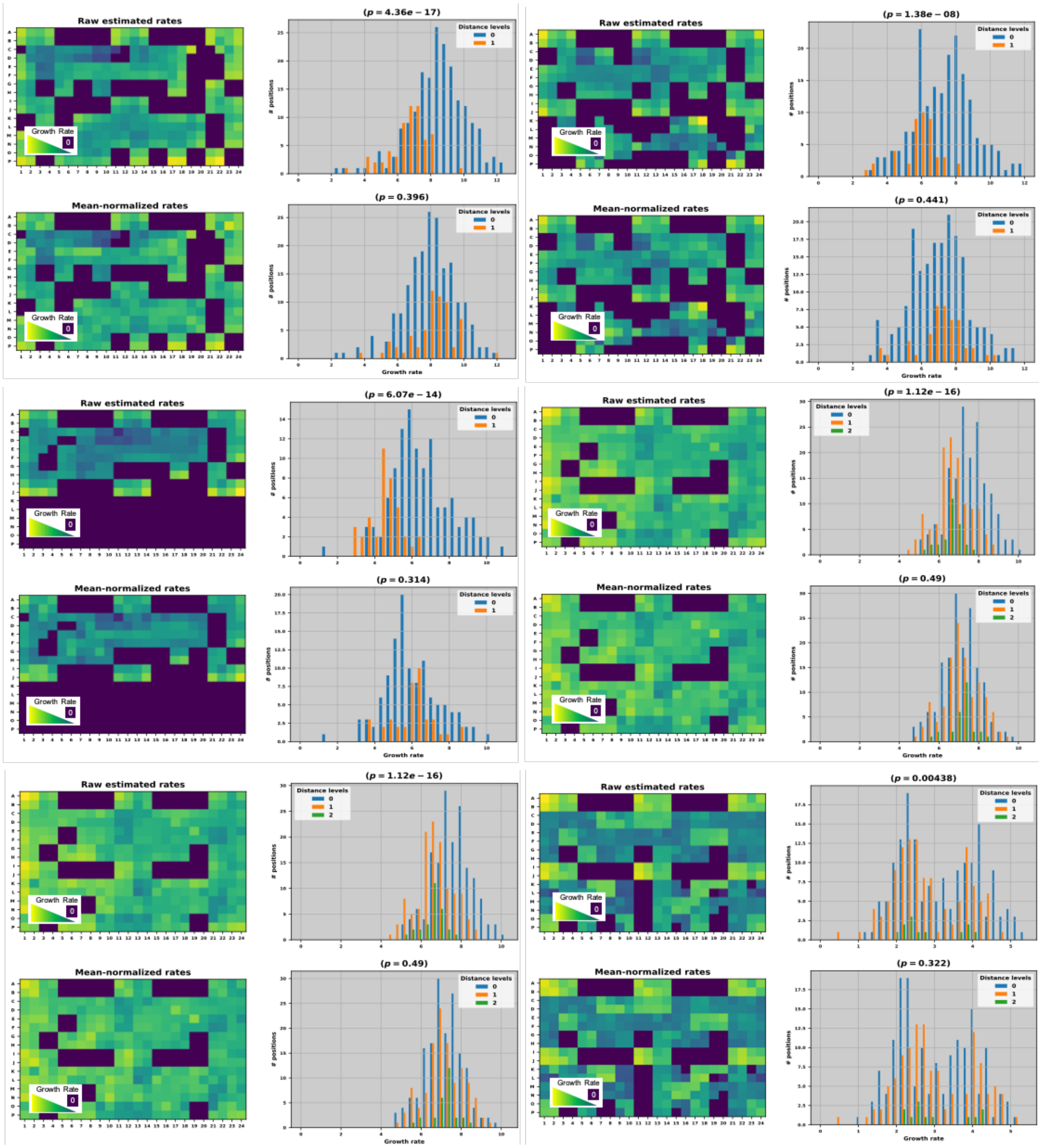


**Figure S2. Normalizations for six representative phenotyping plates.** Normalizations based on the mean growth rate of the wildtype controls on each plate were used to account for edge effects. Heat maps show a visual representation of growth rates across each plate. The X and Y axis are the coordinates for the 384 positions where a colony may appear. In every case, wild-type controls are in rows A, B, I, and J, columns 1-4, 11-14, and 21-24. Histograms compare the growth rate of colonies that are on the edge of the plate or adjacent to an empty position (distance level 0) with those that are one or more positions away from an edge (non-zero distance levels). The p-value reported above the histogram marks the significance of the difference between the growth rate of edge-adjacent colonies and internal colonies. In each case, raw, unnormalized heat maps, histograms, and p-values are shown just above their normalized counterparts.

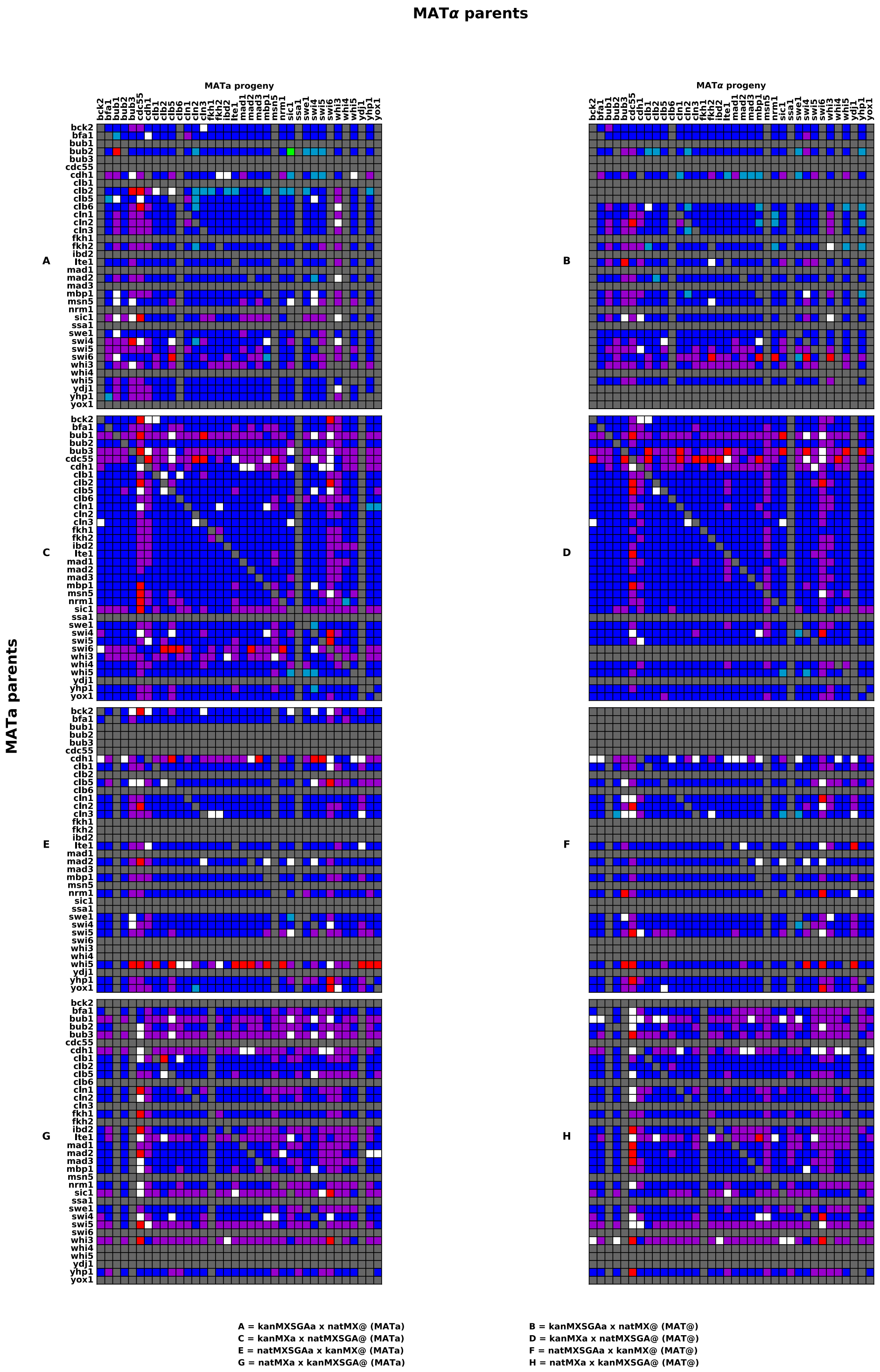


**Figure S3. Comparison of fitness scores for double mutants in all four sets of crosses on YPR media.** White cells indicate zero growth and grey cells indicate missing or excluded data. Royal blue is used to designate fitness scores that differ from WT by fewer than 2 standard deviations. Cyan and green indicate fitness scores that are greater than WT by up to or more than 6 standard deviations respectively. Magenta and red indicate fitness scores that are less than WT by up to or more than 6 standard deviations respectively. **A & B) Cross 1. C & D) Cross 2. E & F) Cross 3. G & H) Cross 4.**

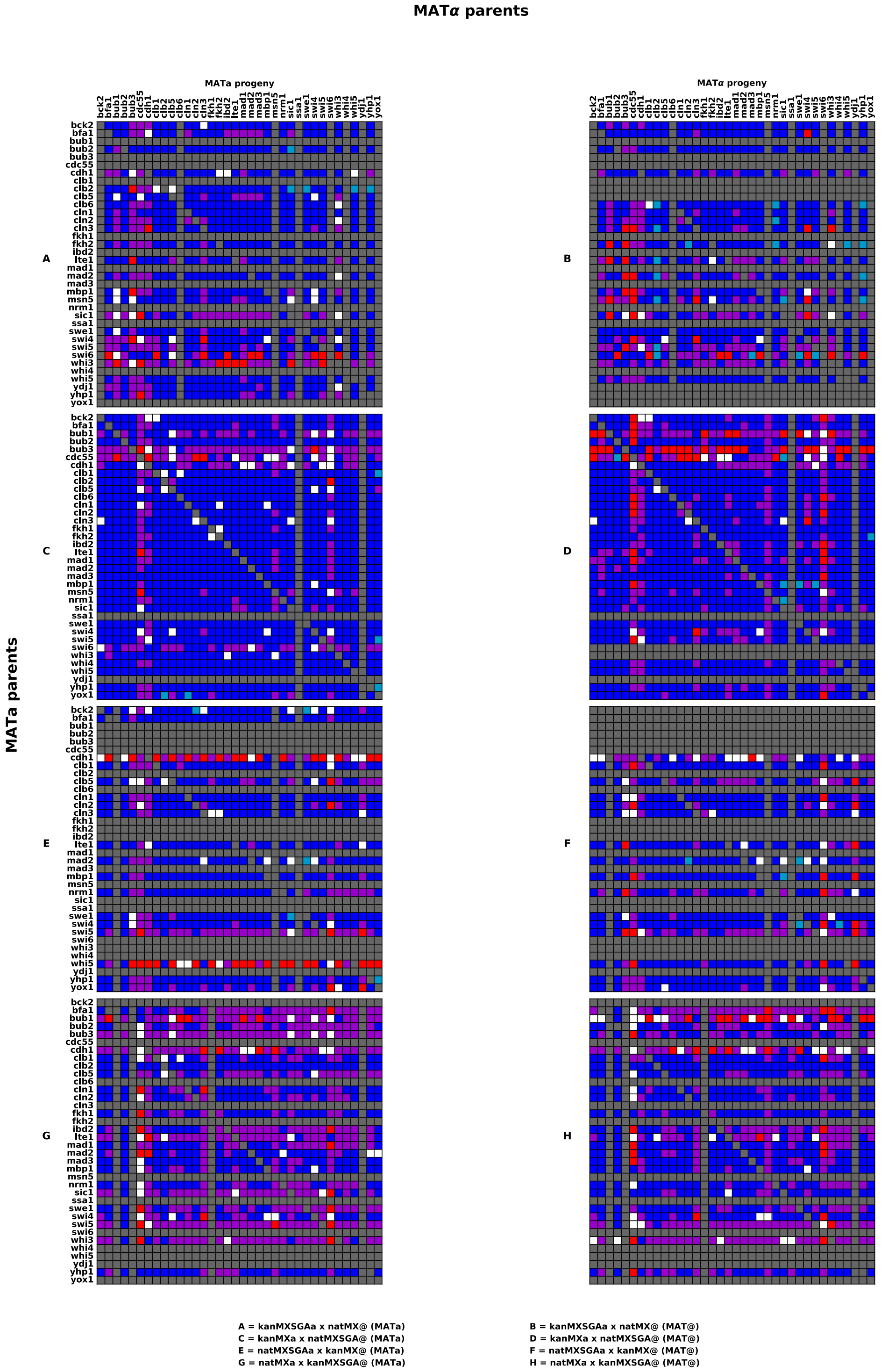


**Figure S4. Comparison of fitness scores for double mutants in all four sets of crosses on YPG media.** White cells indicate zero growth and grey cells indicate missing or excluded data. Royal blue is used to designate fitness scores that differ from WT by fewer than 2 standard deviations.

Cyan and green indicate fitness scores that are greater than WT by up to or more than 6 standard deviations respectively. Magenta and red indicate fitness scores that are less than WT by up to or more than 6 standard deviations respectively. **A & B) Cross 1. C & D) Cross 2. E & F) Cross 3. G & H) Cross 4.**

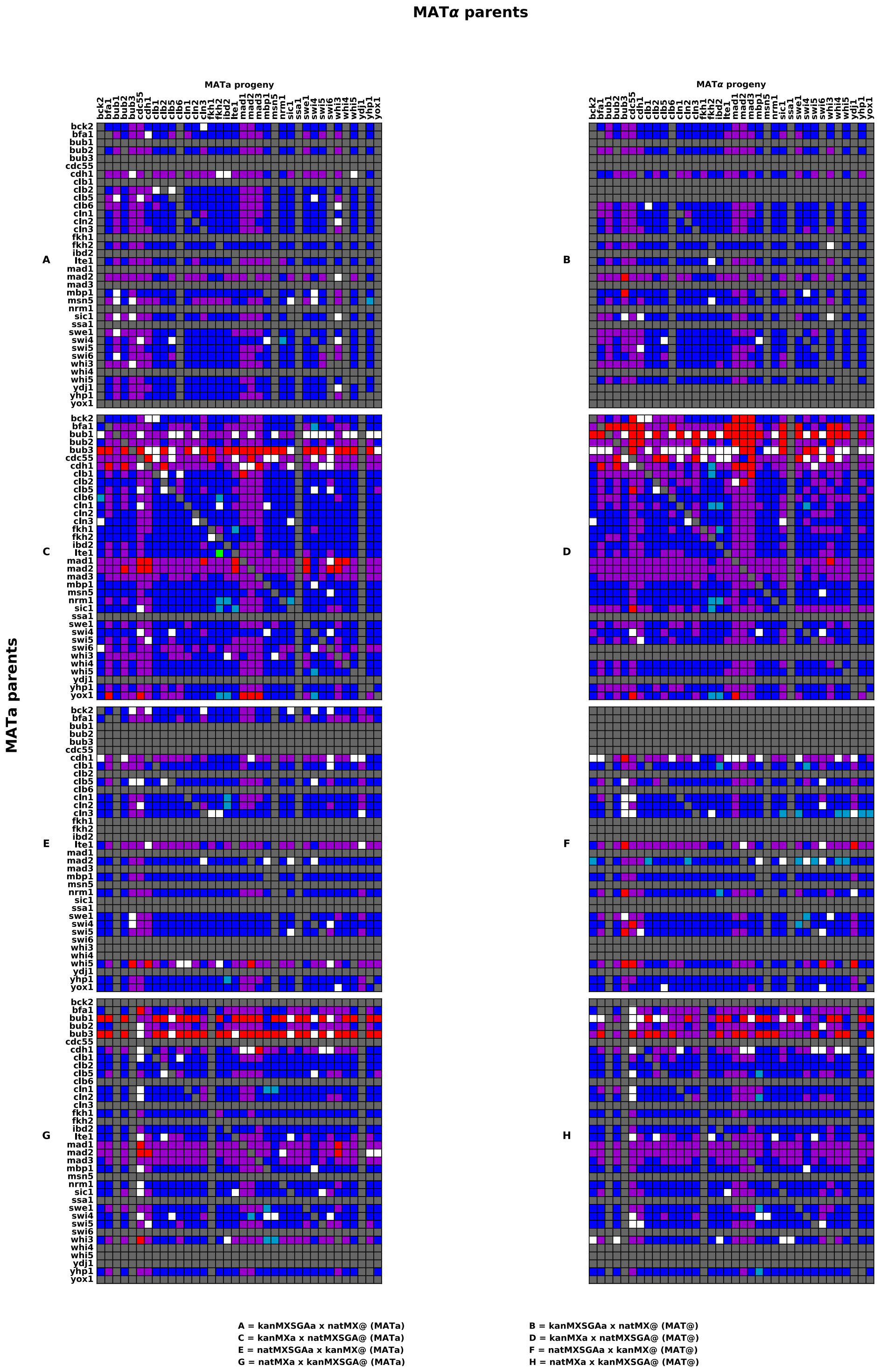


**Figure S5. Comparison of fitness scores for double mutants in all four sets of crosses on YPD-Ben media.** White cells indicate zero growth and grey cells indicate missing or excluded data. Royal blue is used to designate fitness scores that differ from WT by fewer than 2 standard deviations. Cyan and green indicate fitness scores that are greater than WT by up to or more than 6 standard deviations respectively. Magenta and red indicate fitness scores that are less than WT by up to or more than 6 standard deviations respectively. **A & B) Cross 1. C & D) Cross 2. E & F) Cross 3. G & H) Cross 4.**

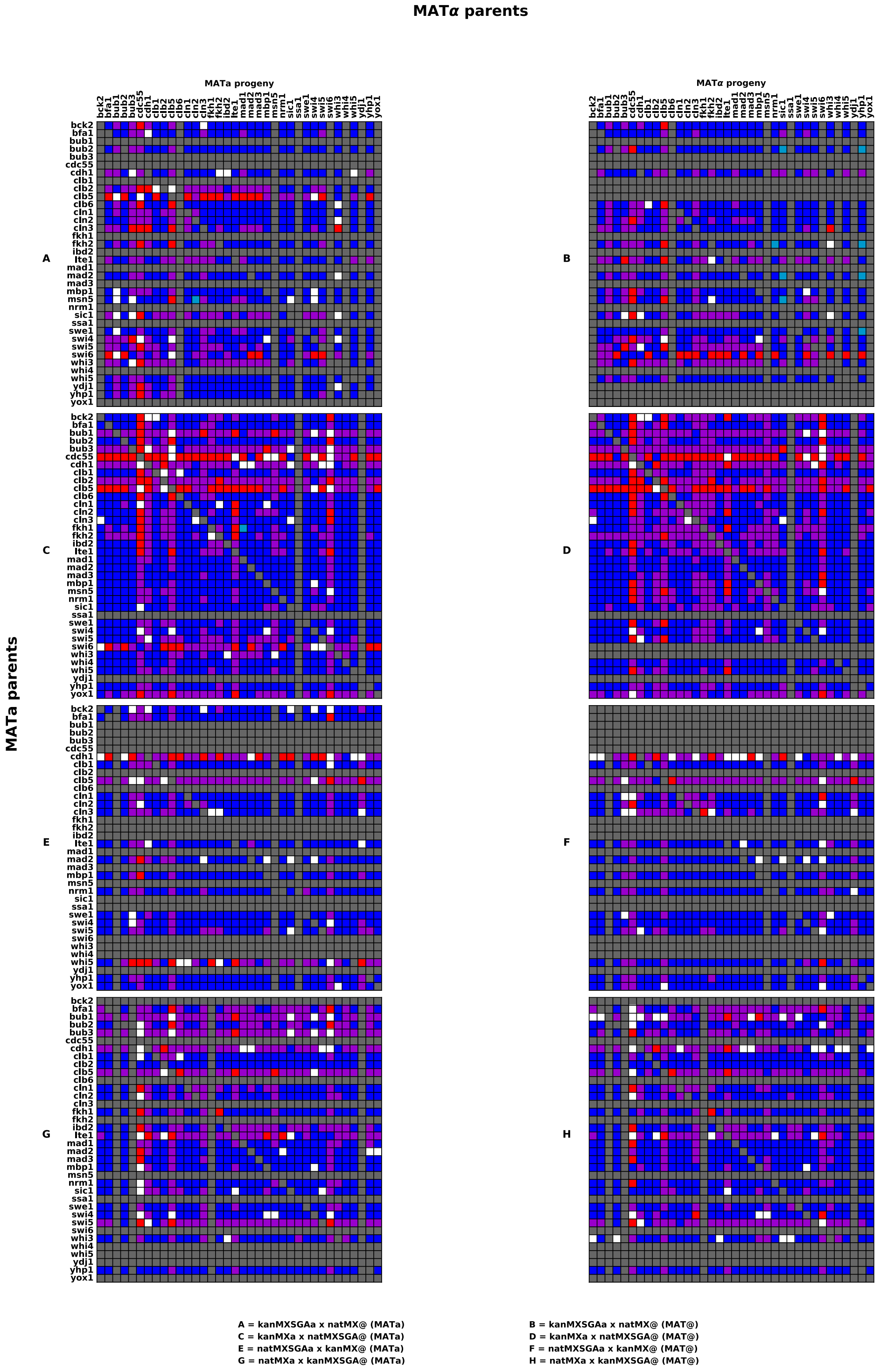


**Figure S6. Comparison of fitness scores for double mutants in all four sets of crosses on YPD-CPT media.** White cells indicate zero growth and grey cells indicate missing or excluded data. Royal blue is used to designate fitness scores that differ from WT by fewer than 2 standard deviations. Cyan and green indicate fitness scores that are greater than WT by up to or more than 6 standard deviations respectively. Magenta and red indicate fitness scores that are less than WT by up to or more than 6 standard deviations respectively. **A & B) Cross 1. C & D) Cross 2. E & F) Cross 3. G & H) Cross 4.**

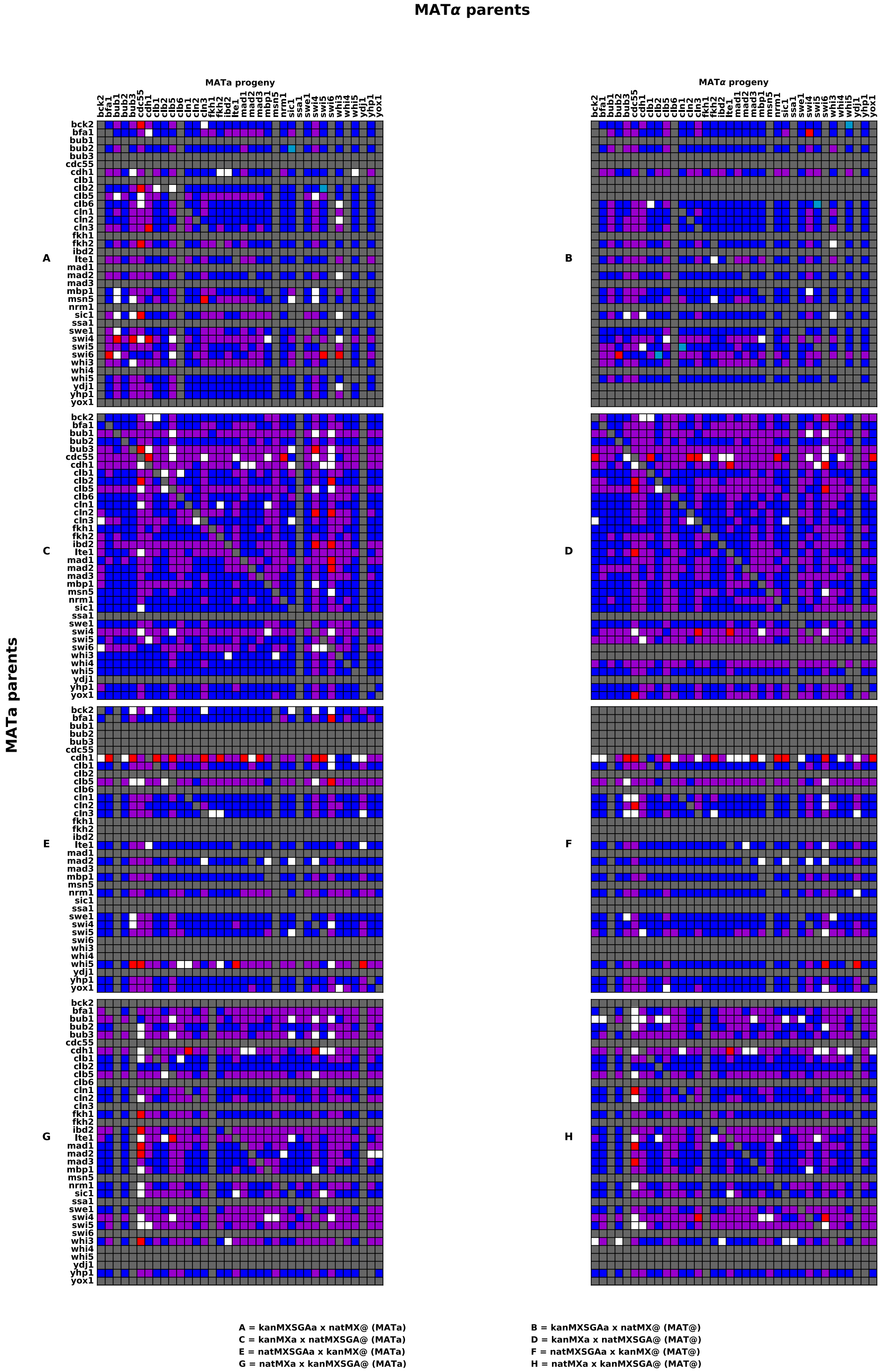


**Figure S7. Comparison of fitness scores for double mutants in all four sets of crosses on YPD-HU media.** White cells indicate zero growth and grey cells indicate missing or excluded data. Royal blue is used to designate fitness scores that differ from WT by fewer than 2 standard deviations. Cyan and green indicate fitness scores that are greater than WT by up to or more than 6 standard deviations respectively. Magenta and red indicate fitness scores that are less than WT by up to or more than 6 standard deviations respectively. **A & B) Cross 1. C & D) Cross 2. E & F) Cross 3. G & H) Cross 4.**

### References


1. Giaever, G. et al. Functional profiling of the Saccharomyces cerevisiae genome. *Nature* **418**, 387-391 (2002).
2. Sherman, F. & Hicks, J. in Guide to yeast genetics and molecular biology, Vol. 194, Edn. 1991/01/01. (ed. C.G.a.G.R. Fink) 21-37 (Academic Press, 1991).
3. Goldstein, A.L. & McCusker, J.H. Three new dominant drug resistance cassettes for gene disruption in Saccharomyces cerevisiae. *Yeast* **15**, 1541-1553 (1999).
4. Knop, M. et al. Epitope tagging of yeast genes using a PCR-based strategy: more tags and improved practical routines. *Yeast* **15**, 963-972 (1999).
5. Longtine, M.S. et al. Additional modules for versatile and economical PCR-based gene deletion and modification in Saccharomyces cerevisiae. *Yeast* **14**, 953-961 (1998).
6. Schiestl, R.H. & Gietz, R.D. High efficiency transformation of intact yeast cells using single stranded nucleic acids as a carrier. *Curr Genet* **16**, 339-346 (1989).
7. Tong, A.H. & Boone, C. Synthetic genetic array analysis in Saccharomyces cerevisiae. *Methods Mol Biol* **313**, 171-192 (2006).
8. OpenWetWare  (OpenWeWare; 2009).
9. Sprague, G.F., Jr. Assay of yeast mating reaction. *Methods Enzymol* **194**, 77-93 (1991).
10. Hsu, W.S. et al. S-phase cyclin-dependent kinases promote sister chromatid cohesion in budding yeast. *Molecular and cellular biology* **31**, 2470-2483 (2011).
11. Stuart, D. & Wittenberg, C. CLB5 and CLB6 are required for premeiotic DNA replication and activation of the meiotic S/M checkpoint. *Genes Dev* **12**, 2698-2710 (1998).
12. Segal, M., Clarke, D.J. & Reed, S.I. Clb5-associated kinase activity is required early in the spindle pathway for correct preanaphase nuclear positioning in Saccharomyces cerevisiae. *The Journal of cell biology* **143**, 135-145 (1998).
13. McCourt, P., Gallo-Ebert, C., Gonghong, Y., Jiang, Y. & Nickels, J.T., Jr. PP2A(Cdc55) regulates G1 cyclin stability. *Cell cycle (Georgetown, Tex.)* **12**, 1201-1210 (2013).
14. Loeb, J.D., Kerentseva, T.A., Pan, T., Sepulveda-Becerra, M. & Liu, H. Saccharomyces cerevisiae G1 cyclins are differentially involved in invasive and pseudohyphal growth independent of the filamentation mitogen-activated protein kinase pathway. *Genetics* **153**, 1535-1546 (1999).
15. Oehlen, L.J. & Cross, F.R. Potential regulation of Ste20 function by the Cln1-Cdc28 and Cln2-Cdc28 cyclin-dependent protein kinases. *J Biol Chem* **273**, 25089-25097 (1998).

16. Levine, K., Huang, K. & Cross, F.R. Saccharomyces cerevisiae G1 cyclins differ in their intrinsic functional specificities. *Molecular and cellular biology* **16**, 6794-6803 (1996).

17. Cross, F.R. Cell cycle arrest caused by CLN gene deficiency in Saccharomyces cerevisiae resembles START-I arrest and is independent of the mating-pheromone signalling pathway. *Molecular and cellular biology* **10**, 6482-6490 (1990).

18. Cvrcková, F. & Nasmyth, K. Yeast G1 cyclins CLN1 and CLN2 and a GAP-like protein have a role in bud formation. *The EMBO journal* **12**, 5277-5286 (1993).

19. Mitchell, D.A. & Sprague, G.F., Jr. The phosphotyrosyl phosphatase activator, Ncs1p (Rrd1p), functions with Cla4p to regulate the G(2)/M transition in Saccharomyces cerevisiae. *Molecular and cellular biology* **21**, 488-500 (2001).

20. Alepuz, P.M., Matheos, D., Cunningham, K.W. & Estruch, F. The Saccharomyces cerevisiae RanGTP-binding protein msn5p is involved in different signal transduction pathways. *Genetics* **153**, 1219-1231 (1999).

21. Pic, A. et al. The forkhead protein Fkh2 is a component of the yeast cell cycle transcription factor SFF. *EMBO J* **19**, 3750-3761 (2000).

22. Becker, J., Walter, W., Yan, W. & Craig, E.A. Functional interaction of cytosolic hsp70 and a DnaJ-related protein, Ydj1p, in protein translocation in vivo. *Molecular and cellular biology* **16**, 4378-4386 (1996).
