## Supplementary figures and images for "Genetic interactions derived from high-throughput phenotyping of 7,350 yeast cell cycle mutants"

### Additional Data Figure 1.tif

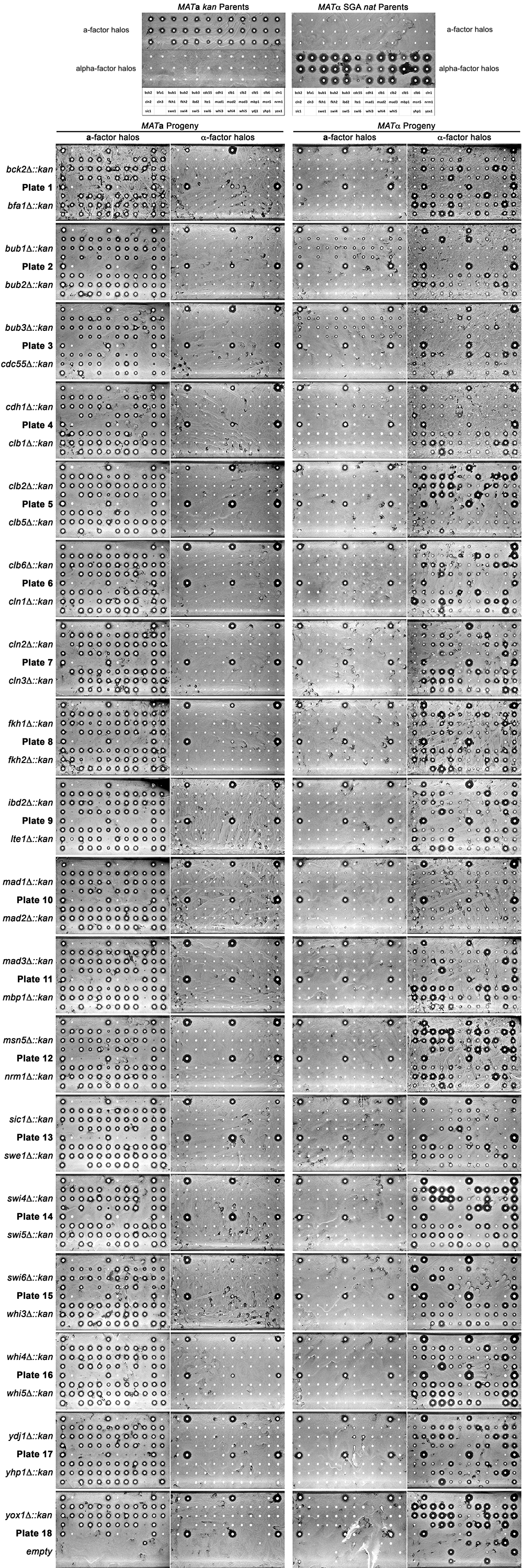

### Additional Data Figure 2.tif

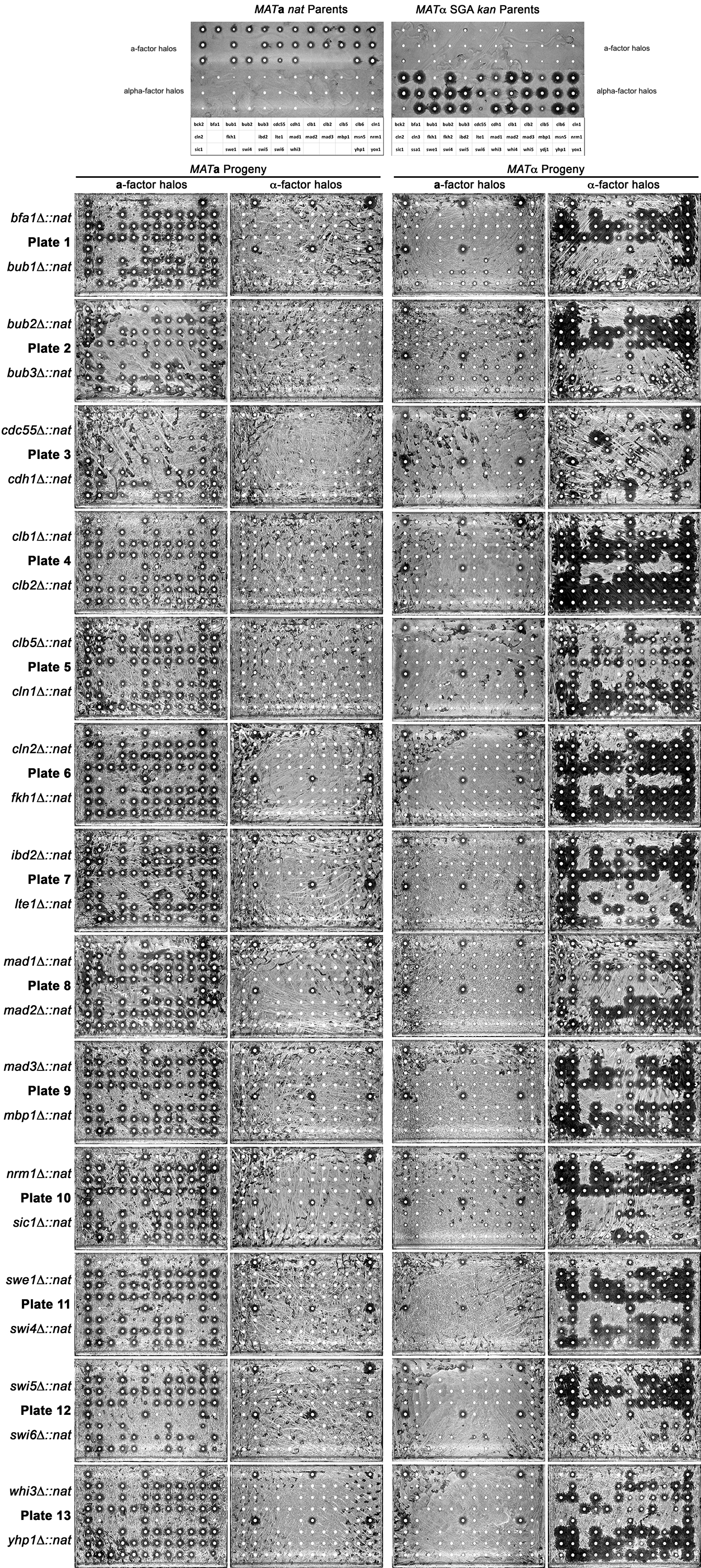

### Additional Data Figure 3.tif

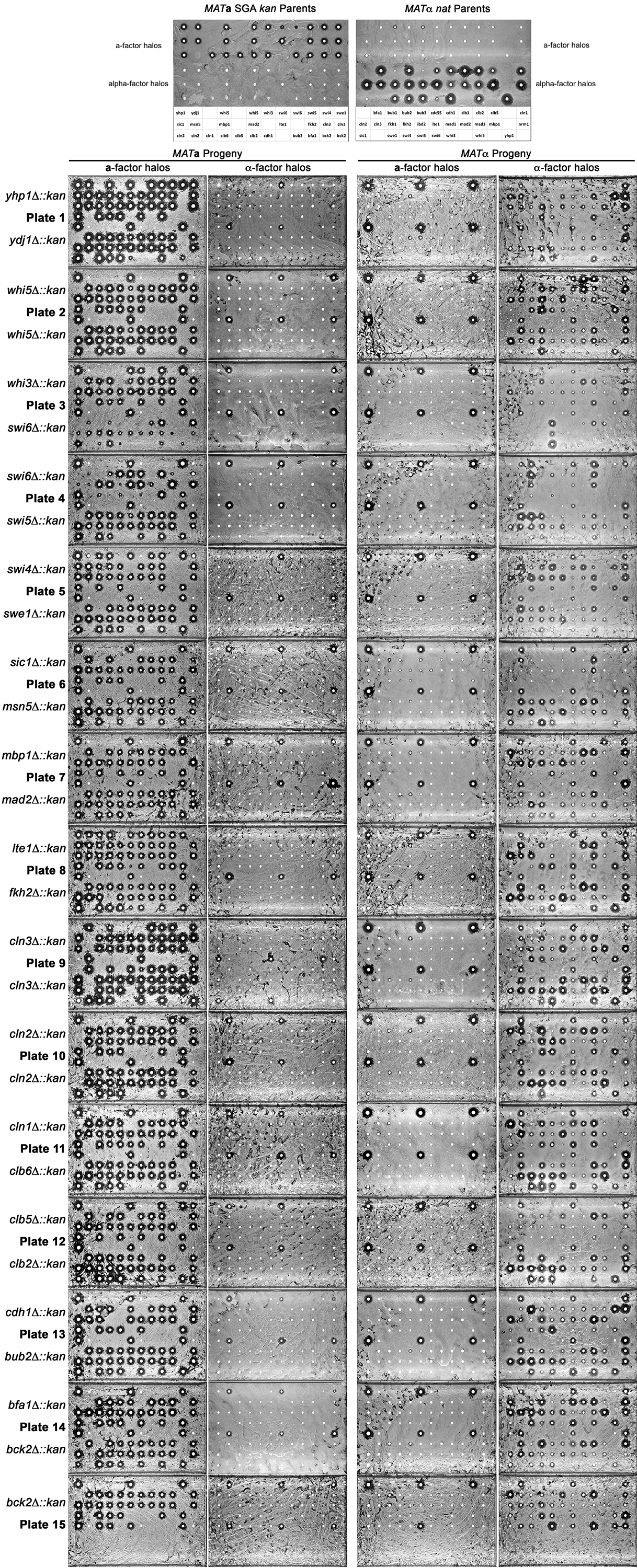

### Additional Data Figure 4.tif

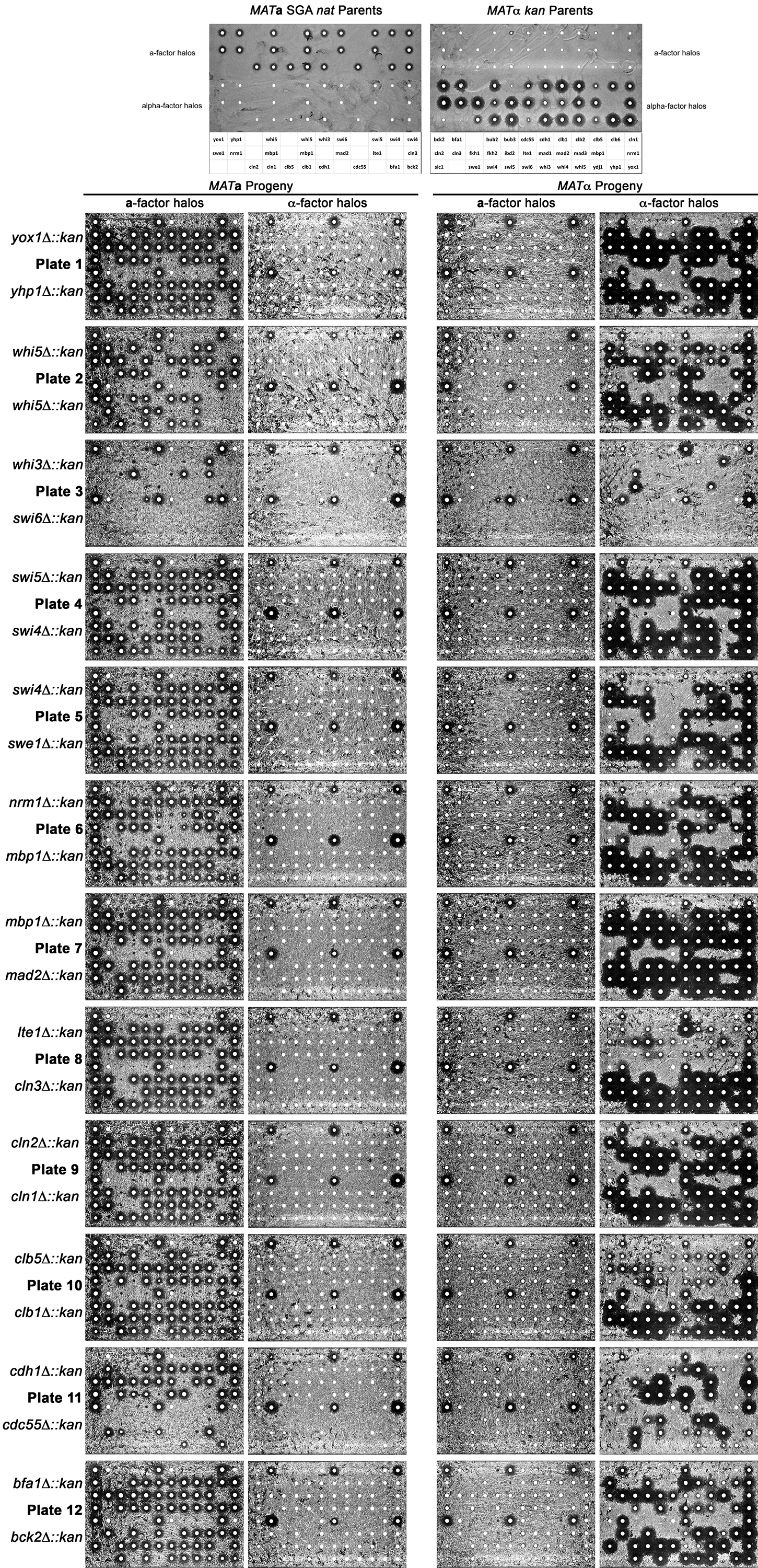

### Additional Data Figure 5.tif

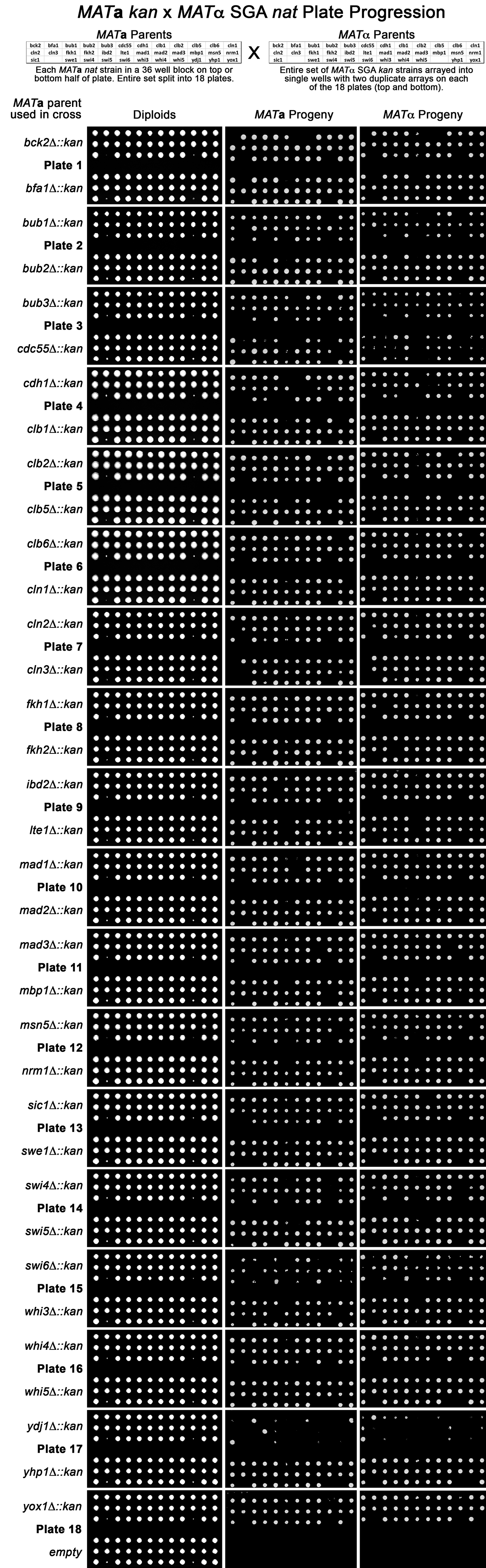

### Additional Data Figure 6.tif

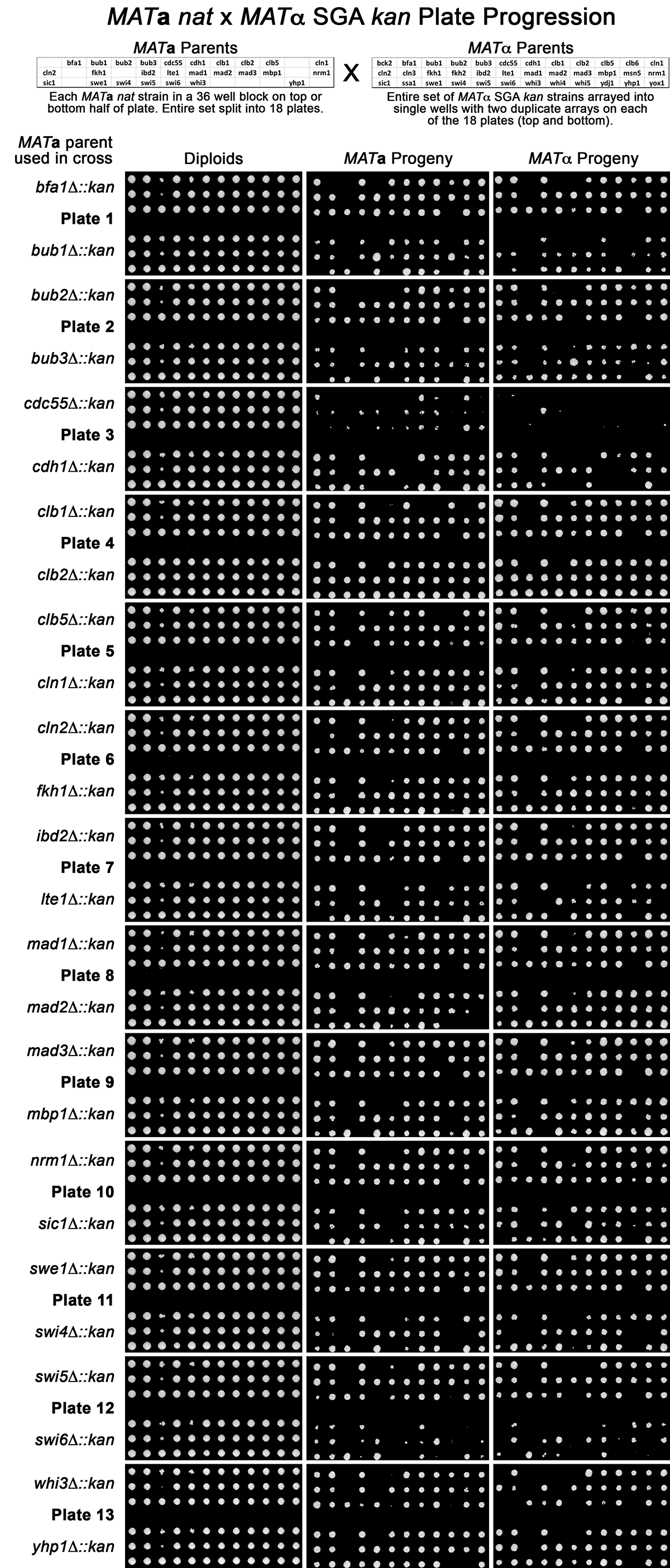

### Additional Data Figure 7.tif

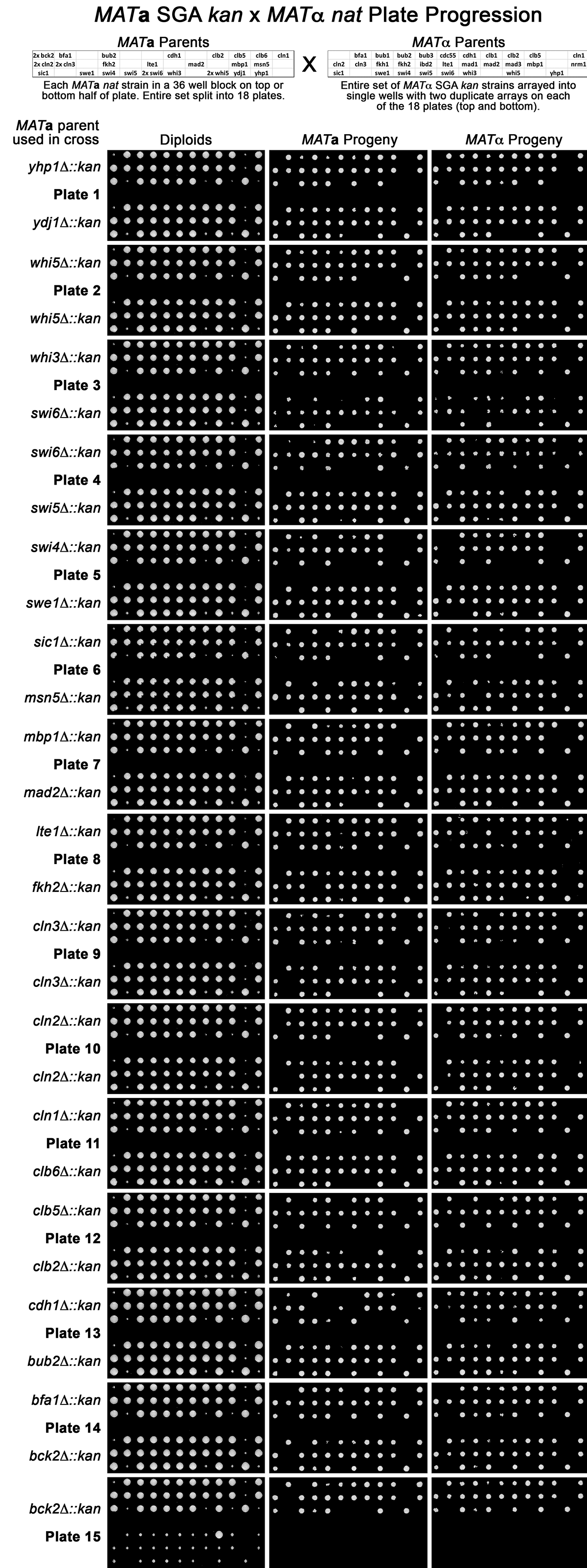

### Additional Data Figure 8.tif

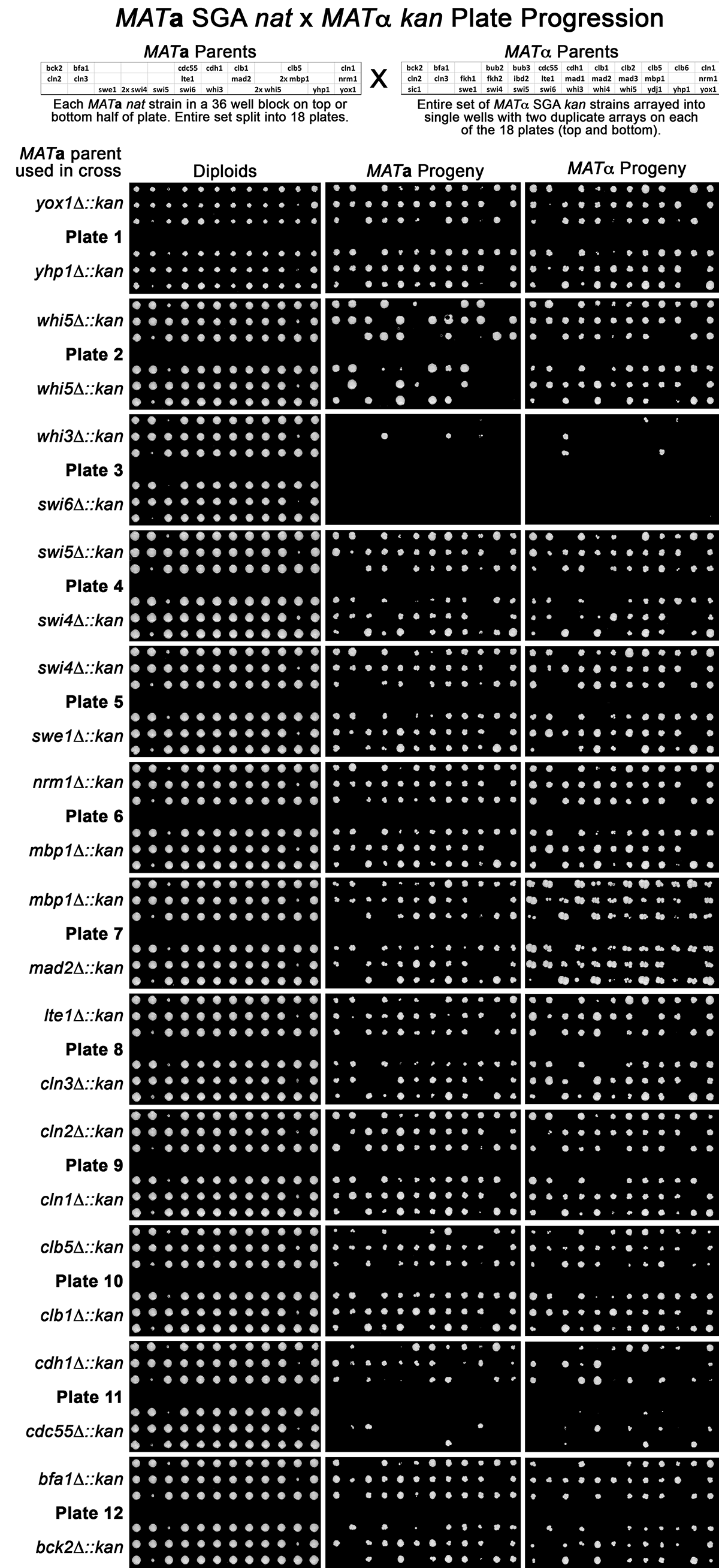
